# Transcription profiles of age-at-maturity-associated genes suggest cell fate commitment regulation as a key factor in the Atlantic salmon maturation process

**DOI:** 10.1101/778498

**Authors:** Johanna Kurko, Paul V. Debes, Andrew House, Tutku Aykanat, Jaakko Erkinaro, Craig R. Primmer

## Abstract

Despite recent taxonomic diversification in studies linking genotype with phenotype, follow-up studies aimed at understanding the molecular processes of such genotype-phenotype associations remain rare. The age at which an individual reaches sexual maturity is an important fitness trait in many wild species. However, the molecular mechanisms regulating maturation timing processes remain obscure. A recent genome-wide association study in Atlantic salmon (*Salmo salar*) identified large-effect age-at-maturity-associated chromosomal regions including genes *vgll3*, *akap11* and *six6*, which have roles in adipogenesis, spermatogenesis and the hypothalamic-pituitary-gonadal (HPG) axis, respectively. Here, we determine expression patterns of these genes during salmon development and their potential molecular partners and pathways. Using Nanostring transcription profiling technology, we show development- and tissue-specific mRNA expression patterns for *vgll3*, *akap11* and *six6*. Correlated expression levels of *vgll3* and *akap11*, which have adjacent chromosomal location, suggests they may have shared regulation. Further, *vgll3* correlating with *arhgap6* and *yap1*, and *akap11* with *lats1* and *yap1* suggests that Vgll3 and Akap11 take part in actin cytoskeleton regulation. Tissue-specific expression results indicate that *vgll3* and *akap11* paralogs have sex-dependent expression patterns in gonads. Moreover, *six6* correlating with *slc38a6* and *rtn1*, and Hippo signaling genes suggests that Six6 could have a broader role in the HPG neuroendrocrine and cell fate commitment regulation, respectively. We conclude that Vgll3, Akap11 and Six6 may influence Atlantic salmon maturation timing via affecting on adipogenesis and gametogenesis by regulating cell fate commitment and the HPG axis. These results may help to unravel general molecular mechanisms behind maturation.

## Introduction

One of the classic challenges in biological research, linking genotype with phenotype, has seen a dramatic taxonomic diversification in recent years as new genomic technologies have enabled genomic approaches to be conducted in almost any species. However, such diversification is not yet apparent when it comes to understanding the molecular processes by which genotypes are translated to ecologically-relevant phenotypes despite the fundamental and applied significance (1–3). Age-at-maturity is closely linked to fitness in many species, with the timing of maturation often involving trade-offs in reproductive strategies shaping maturation timing variation (4). For example, delayed maturation can lead to larger body size, higher fecundity and increased offspring survival, but longer generation times can carry an increased mortality risk prior to reproduction by prolonging pre-maturity life stages (5). A recent genome-wide association study (GWAS) in Atlantic salmon (*Salmo salar*) identified a single locus on chromosome 25 that associates strongly with age-at-maturity, and a single nucleotide polymorphism (SNP) located near the gene *vgll3* (vestigial-like family member 3) explained as much as 39 % of the phenotypic variation in maturation age in both males and females (6, 7). In addition to Atlantic salmon, *vgll3* has also been linked with pubertal timing, growth and body condition in humans (8–10), which indicates that it may have an evolutionarily conserved role in the regulation of vertebrate maturation timing, the trait generally being polygenic (8, 9, 11–13). The identification of a large-effect locus in salmon provides a rare opportunity to investigate the molecular processes behind this association.

Sexual maturation is a biological process stemming from a complex chain of events culminating in the first reproduction. The maturation process commences already in the embryo after fertilization by allocating energy to the growth and differentiation of developing gonads and is completed when gametes are produced (14–16). Although timing of maturation is known to be mediated by interplay between fat accumulation and activation of the hypothalamic-pituitary-gonadal (HPG) axis (17–19) the exact molecular mechanisms regulating the process are still obscure.

Maturation requires sufficient fat storage to provide energy for proper gonad development. Therefore, evidence showing that *vgll3* encodes a negative regulator of adipocyte maturation and that its mRNA expression inversely correlates with total body weight and fat content in mice (20), suggests *vgll3* is a good candidate for having a role in sexual maturation in salmon. A recent study linking *vgll3* with reduced adiposity indices in the Mongolian human population provides further evidence for general, species-wide role of *vgll3* in adipose regulation (21). Beyond regulating adipocyte differentiation, Vgll3 has also been shown to have a broader role in mesenchymal-derived cell fate decision. Studies show that Vgll3 promotes expression of the chondrocyte and osteocyte inducing markers in murine preadipocyte cell line (20) and myogenesis in mouse and human myoblasts (22). Consistently, *vgll3* overexpression is found in sarcomas (23) and cartilage with endemic osteoarthritis (24). During embryonic development (25–27) and in adult vertebrates (22, 25, 28, 29), the expression of *vgll3* has been detected in various tissues, including skeletal muscle, skin, heart, lung, gill, nose, kidney, liver, spleen, stomach, gut, pyloric caeca, brain and eye, potentially suggesting a broad role in development. *vgll3* expression is also observed in testis (25, 29, 30) and ovary (29, 31), further supporting the participation of Vgll3 in sexual maturation. The exact molecular mechanisms via which Vgll3 operates on cell fate determination, and also maturation, are unclear, but it is known to be a cofactor for all known Tead transcription factors (22, 27). By binding to Teads, Vgll3 has been shown to influence the Hippo signaling pathway (22) that regulates cell fate commitment and organ growth (32, 33). In line with cell fate regulation, it has been demonstrated earlier that Vgll3 may also have a role in tumor suppression in ovarian cancer (31). Furthermore, one of the Tead genes, *tead3*, and other major Hippo pathway members have been linked with maturation in Atlantic salmon (29, 34) and other species (35–37) emphasizing the role of the Hippo pathway in the maturation process.

In addition to *vgll3*, two other genes, *akap11* (on chromosome 25) and *six6* (on chromosome 9), associate with age-at-maturity in Atlantic salmon (6). However, association of *six6* with maturation timing is only seen before population structure correction. In addition to salmon, *SIX6* (SIX homeobox 6) associates with age-at-menarche and adult height in humans (11) and puberty in cattle (38), and its role in reproduction has been widely studied in several vertebrates. *Six6* encodes a transcription factor which is detected early in the development in the anterior neural plate in regions that comprises the eye field and prospective hypothalamus, pituitary gland and olfactory placodes (39). Later in development, *six6* expression is seen in the hypothalamus, pituitary gland and testis, organs of the HPG axis (39–43). Accordingly, studies in mice show that Six6 is required for fertility by regulating the maturation of gonadotropin-releasing hormone (GnRH) neurons and expression of GnRH in the hypothalamus (44) and repressing transcription of the β-subunit genes of gonadotropin hormones in pituitary gonadotrope cell line (43). In addition to being an important regulator of the HPG axis, Six6 has an essential role in eye development (39–42), for example controlling photoreceptor differentiation (45), which is crucial for photoperiod sensing in seasonal breeders such as salmon (46). Contrary to *six6* whose role in testis remains so far unknown, *akap11*, although expressed in most studied tissues and cell types, has the strongest signal and suggested role in testis and sperm (47). It encodes A-kinase anchor protein 11, which interacts with type I and II regulatory subunits of protein kinase A (PKA) in order to tether PKA to discrete locations within a cell to control many essential functions, such as cell cycle (48) and lipid metabolism (49). Expression of Akap11 mRNA and protein is observed in sperm throughout spermatogenesis and its suggested association with cytoskeletal structure indicates its importance in the sperm function (47), and thus maturation. Although there is information available about the expression patterns and molecular functions of the age-at-maturity-associated genes *vgll3*, *akap11* and *six6* in some vertebrate species, studies covering a range of developmental time points are lacking. Therefore, in order to be able to determine functional molecular mechanisms of these genes during maturation, first, it is crucial to know when and where these genes are expressed, and what are the potential molecular partners and pathways they link with. Since Atlantic salmon has an exceptionally large-effect locus associating with age-at-maturity, it provides an attractive model for studying molecular mechanisms linked with sexual maturation. Thus, our aim was to investigate the expression patterns of *vgll3*, *akap11* and *six6*, and additional known key genes related to their suggested functions and pathways in a range of Atlantic salmon developmental time points and tissues. Using Nanostring technology, we characterized expression profiles of *vgll3*, *akap11* and *six6* paralogs and 205 other genes related to their functions in five Atlantic salmon developmental stages and 15 tissues. Based on our results, we hypothesize a novel role for Vgll3 in participating in cell fate control via actin cytoskeleton regulation, and Akap11 assisting in this process. In addition, we suggest that Six6 may associate broadly with neuroendocrine secretion regulation in the HPG axis, and have a direct role in testis function.

## Materials and Methods

### Animals

The Atlantic salmon (*Salmo salar*) used in this study were derived from hatchery-maintained Neva river strain at the Natural Resources Institute Finland (62°24′50″N, 025°57′15″E, Laukaa, Finland) and the Inarijoki river, headwater river of the Teno river, near Angeli in northern Finland. Three-year-old wild parr (freshwater stage individuals) from the Inarijoki were caught by electrofishing in early September 2016. The hatchery-maintained Neva strain eggs were fertilized in November 2016 and incubated in circulated water system until hatching. After hatching, alevin (hatched individuals) and fry (individuals able to feed) were grown in standard commercial fish farming conditions. After 14 days from hatching, some alevin were transferred to be grown in hatchery at the Lammi Biological Station (61°04′45″N, 025°00′40″E, Lammi, Finland). Relative ages of the embryos, alevin and fry were calculated in degree days (°d) (number of days after fertilization multiplied by temperature in °C), which are used throughout the text for the embryonic stages. In addition, τ_s_ units (time it takes an embryo to form one somite pair in certain temperature in °C) were calculated according to Gorodilov (50). Three-year-old hatchery-maintained Neva river parr were reared in standard commercial fish farming conditions at the Natural Resources Institute Finland hatchery in Laukaa, Finland.

### Sample collections

Pre-hatched embryos from two developmental time points (186 °d/177 τ_s_ and 258 °d/260 τ_s_) were dissected from eggs, and alevin (436 °d/377 τ_s_) and fry (1119 °d, 1320 °d and 2212 °d) were caught by a plastic Pasteur pipette and net, respectively. From the embryos and alevin, yolk sac was excised with a scalpel. All samples were stored in RNA preservation buffer (25 mM sodium citrate, 10 mM EDTA, 70 g ammonium sulfate/100 ml solution, pH 5.2) at −20 °C, whereby fry were first euthanized by anesthetic overdose of Tricaine methanesulfonate. Of these, four 186 °d embryos, four 258 °d embryos, four alevin, two normal diet fed 1119 °d fry, all from Natural Resources Institute Finland, and four 1320 °d fry, of which two were fed with normal and two with low-fat diet, from Lammi Biological Station, were chosen for further analysis. In addition, blood samples were extracted from the caudal vein of two normal diet fed and two low-fat diet fed 2212 °d fry from Lammi Biological Station and stored in RNAprotect Animal Blood Tubes (Qiagen, Hilden, Germany) at −20 °C. Samples from adipose, brain, eye, fin (adipose and caudal), gill, gonad, heart, kidney, liver, muscle, pyloric caeca, spleen and skin tissues were dissected from four hatchery-maintained Neva river parr (two males and two females) and four wild Inarijoki parr (two immature males and two mature males) and stored in RNA preservation buffer at −20 °C. As not all of the above-mentioned tissues were collected from all eight fish, the specific tissue samples assessed for each individual in this study are described in Additional file 1.

### RNA extraction

Altogether 96 samples, including whole embryos, alevin and fry, and tissue samples from parr, were disrupted and homogenized in the presence of NucleoZOL (Macherey-Nagel, Düren, Germany) for lysis in 2 ml safe-lock tubes containing one 5 mm stainless steel bead (Qiagen) for 2.5-3 minutes at 30 Hz using TissueLyzer II (Qiagen). Total RNA was extracted from the lysate using NucleoSpin RNA Set for NucleoZOL (Macherey-Nagel) according to the manufacturer’s instructions. After the elution step, RNA was treated with rDNase using the rDNase Set (Macherey-Nagel) to remove any residual DNA and, subsequently, purified with either NucleoSpin RNA clean-up or NucleoSpin RNA clean-up XS kit(Macherey-Nagel) according to the RNA yield. Blood samples were lysed in the RNAprotect Animal Blood Tubes (Qiagen) and total RNA was extracted using the RNeasy Protect Animal Blood System kit (Qiagen) according to the manufacturer’s protocol followed by concentration of RNA using NucleoSpin RNA clean-up XS kit (Macherey-Nagel). The quantity and quality of RNA was determined using both NanoDrop Spectrophotometer ND-1000 (Thermo Scientific, Wilmington, DE, USA) and 2100 BioAnalyzer system (Agilent Technologies, Santa Clara, CA, USA).

### Nanostring mRNA expression panel

A total of 220 genes were chosen for analysis based on information from the literature, the IPA (Ingenuity Pathway Analysis) tool (Qiagen) and other freely available web tools and databases. These included the age-at-maturity-associated genes *vgll3a* and *akap11a* on chromosome 25, and *six6a* on chromosome 9, and their corresponding paralogs *vgll3b* and *akap11b* on chromosome 21, and *six6b* on chromosome 1, as well as 205 genes potentially linked to these age-at-maturity-associated genes based on their functions and pathways. Further, nine commonly used potential housekeeping genes (*actb*, *ef1aa*, *ef1ab*, *ef1ac*, *gapdh*, *hprt1*, *rpabc2a*, *rpabc2b* and *rps20*) were included in the gene panel as candidates for data normalization. Because of the duplicated Atlantic salmon genome, most genes possess one or more paralogs in their counterpart chromosomes. Therefore, all paralogs of each gene of interest were searched using the SalmoBase (http://salmobase.org/) and NCBI RefSeq databases and included in this study. The exception was that for the genes surrounding the age-at-maturity-associated genes in the chromosomes 25 and 9 according to (6) the possible paralogs were excluded, as those particular genes were only of interest. Paralogs were renamed by adding a letter at the end of their name alphabetically in order to separate them as shown above. Gene accession numbers, IDs, full names and functional categories are listed in Additional file 2. Genes were grouped into three different functional categories, “Cell fate”, “Metabolism” and “HPG axis”, based on their known functions and pathways. Many of the genes have several functions and could thus be placed in multiple categories whereby the chosen categories were the most relevant to the current study. In addition, genes in the chromosomes 25 and 9 were included in the “Chr 25 candidate region” and “Chr 9 candidate region” categories, respectively. mRNA expression levels of chosen genes were studied using Nanostring technology (NanoString Technologies, Seattle, WA, USA). Probes for each gene paralog, targeted at all known transcript variants, were designed using reference sequences in the NCBI RefSeq database. For some paralogs, it was impossible to design specific probes, as sequence similarity between paralogs was too high. Altogether, 96 samples were analyzed using nCounter Custom CodeSet for probes targeting 220 genes and nCounter Master kit (NanoString Technologies). First, 100 ng of the RNA of each sample was denatured, after which the probes were hybridized with the RNA overnight. Post-hybridization purification and image scanning was performed the following day.

### Data analysis

Six genes (*ef1ac, prkar1aa, rps20, rxrbaa, vdraa* and *vdrab*) (Additional file 2) were selected from the 220 genes in the gene panel for use in data normalization as they exhibited a low coefficient of variation and average count values across samples. These genes included two (*ef1ac* and *rps20*) of the nine genes originally included as potential normalization candidates, as well as four additional genes (*prkar1aa, rxrbaa, vdraa* and *vdrab*). Raw count data from the Nanostring mRNA expression analysis was normalized by RNA content normalization factor for each sample calculated from geometric mean count values of these six genes. After normalization and quality control, two samples (shown in the Additional file 1) were removed from the data due to too high RNA content normalization factor values (> 3.0). Quality control and normalization of the data was performed using the nSolver Analysis Software (v3.0) (NanoString Technologies). Mean count values of genes of interest were calculated for all four developmental stages and 15 tissues. For fin, mean count value was averaged across adipose and tail fin samples (no difference was apparent, see results). A normalized count value of 20 was set as a background signal threshold. Forty-two genes were on average below background signal across all stages, which left 178 genes to estimate 507 pairwise correlations (plus one extra, see below) with the age-at-maturity-associated genes. To estimate 507 pairwise correlations between the three age-at-maturity-associated genes and 178 of the 211 studied genes with average expression above the threshold across the early developmental stages, correlation coefficients were determined among residuals of normalized count data that were controlled for different means of genes and developmental stages, using bivariate linear models under residual maximum likelihood. Correlation standard errors were approximated using a Taylor series. To determine significance, models with unstructured covariance vs. diagonal residual covariance structure were compared using likelihood ratio tests (LRT) (51). As this was an exploratory study, *P* < 0.01 was considered relevant in LRTs. Genes with expression levels correlating with the age-at-maturity-associated genes were included in analyzing temporal transcript variation during early development. Specifically, a linear mixed model was fitted under residual maximum likelihood for normalized count data with a random sample term to account for technical among-sample variation and with fixed effects for 24 retained genes (see results), four stages and their interaction. The genes exhibited different variances (LRT between model with homo- vs. heteroscedastic residual variance; 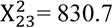, *P* < 0.001) and residual variance was, therefore, allowed to be heteroscedastic. Model terms were tested using Wald’s *F*-test and gene-specific pairwise comparisons among the four early developmental stages were conducted, whereby *p*-values were adjusted for the false discovery rate (*FDR*) according to (52) and comparisons with *FDR* < 0.05 were regarded as relevant. Linear models were estimated using ASReml-R (53) and data analysis was performed using the R statistical software. The custom R script used to analyze the data is shown in Additional file 3.

### Data Availability Statement

**Additional file 1.** Tissues of eight fish chosen for the study.

**Additional file 2.** Table of genes included in the study.

**Additional file 3.** Custom R script used to analyze the data.

**Raw data.** mRNA expression data for genes reported in the study will be submitted to NCBI Gene Expression Omnibus upon acceptance of the study

## Results

### *Vgll3*, *akap11* and *six6* expression during early developmental stages

mRNA expression levels of age-at-maturity-associated genes and their paralogs were studied at four early Atlantic salmon developmental stages (186 °d and 258 °d embryos, alevin and fry). *Vgll3a* was expressed at low levels during all four developmental stages with the highest expression in alevin (Fig. 1). *Six6a* expression level was also overall low and declined from the embryonic stages towards fry (Fig. 1). Both *akap11* paralogs were expressed at low levels in all stages although *akap11b* had clearly higher expression throughout development than *akap11a* (Fig. 1). On the contrary, expression of *vgll3b* and *six6b* paralogs that do not associate with age-at-maturity was below background count level during all studied developmental stages (Fig. 1). This indicates that the paralogs associated with maturation timing are expressed throughout early salmon development.

**Fig. 1.**
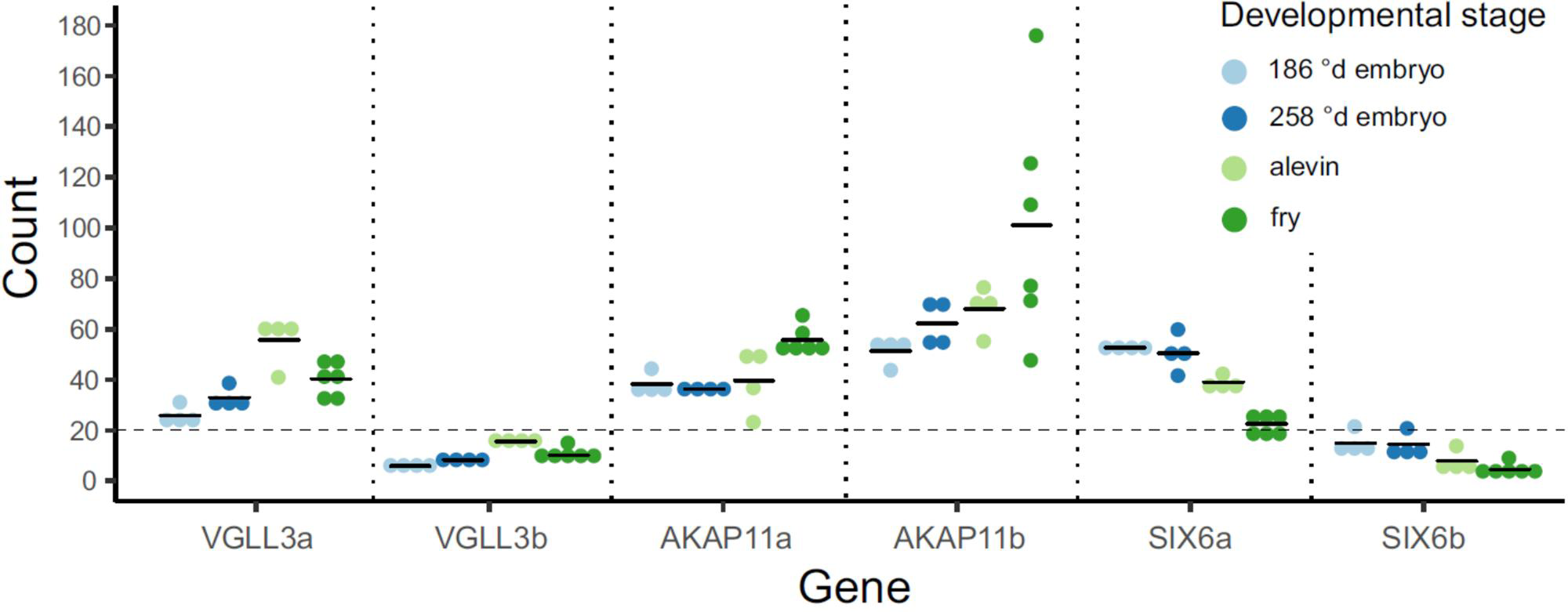
mRNA expression patterns of the age-at-maturity-associated genes *vgll3a*, *akap11a* and *six6a*, and their paralogs *vgll3b, akap11b* and *six6b* during early Atlantic salmon developmental stages. Twenty was set as a count baseline (dashed line) above which counts were considered as detected. The data are represented as individual (dot) and mean (line) counts.

### Genes with expression patterns correlating with *vgll3a*, *akap11a* and *six6a*

In order to identify genes that may be involved in the same molecular networks along with the age-at-maturity-associated genes *vgll3a*, *akap11a* and *six6a*, we determined the expression levels of 205 additional genes (Additional file 2) previously shown to have related functions or suggested as members of pathways linked with puberty and adiposity (see Methods). Pairwise correlations of these genes with *vgll3a*, *akap11a* and *six6a* were then estimated across the expression levels in samples from the four above-mentioned early developmental stages. Models for seven correlations did not converge due to singularities. Twenty-six correlations yielded results with *P* < 0.01 (range = 0.0001-0.009). Interestingly, expression of *vgll3a* correlated positively with one of the other age-at-maturity genes, *akap11a*, as well as with *arhgap6e* and negatively with *rd3l* and *yap1* (Fig. 2)*. Akap11a* correlated positively, in addition to *vgll3a*, with *nr1i2* and negatively with *foxp3a*, *lats1a*, *sox9d*, *rd3l* and *yap1* (Fig. 2). Because *rd3l* and *yap1* correlated with both *vgll3a* and *akap11a*, the correlation between the two was also estimated, and, indeed, a positive correlation was detected between them (Fig. 2). *Six6a* correlated positively with twelve genes related to, for example, the HPG axis signaling, eye development and PKA function, and negatively with *vdrab* and *egr1d* (Fig. 2).

**Fig. 2.**
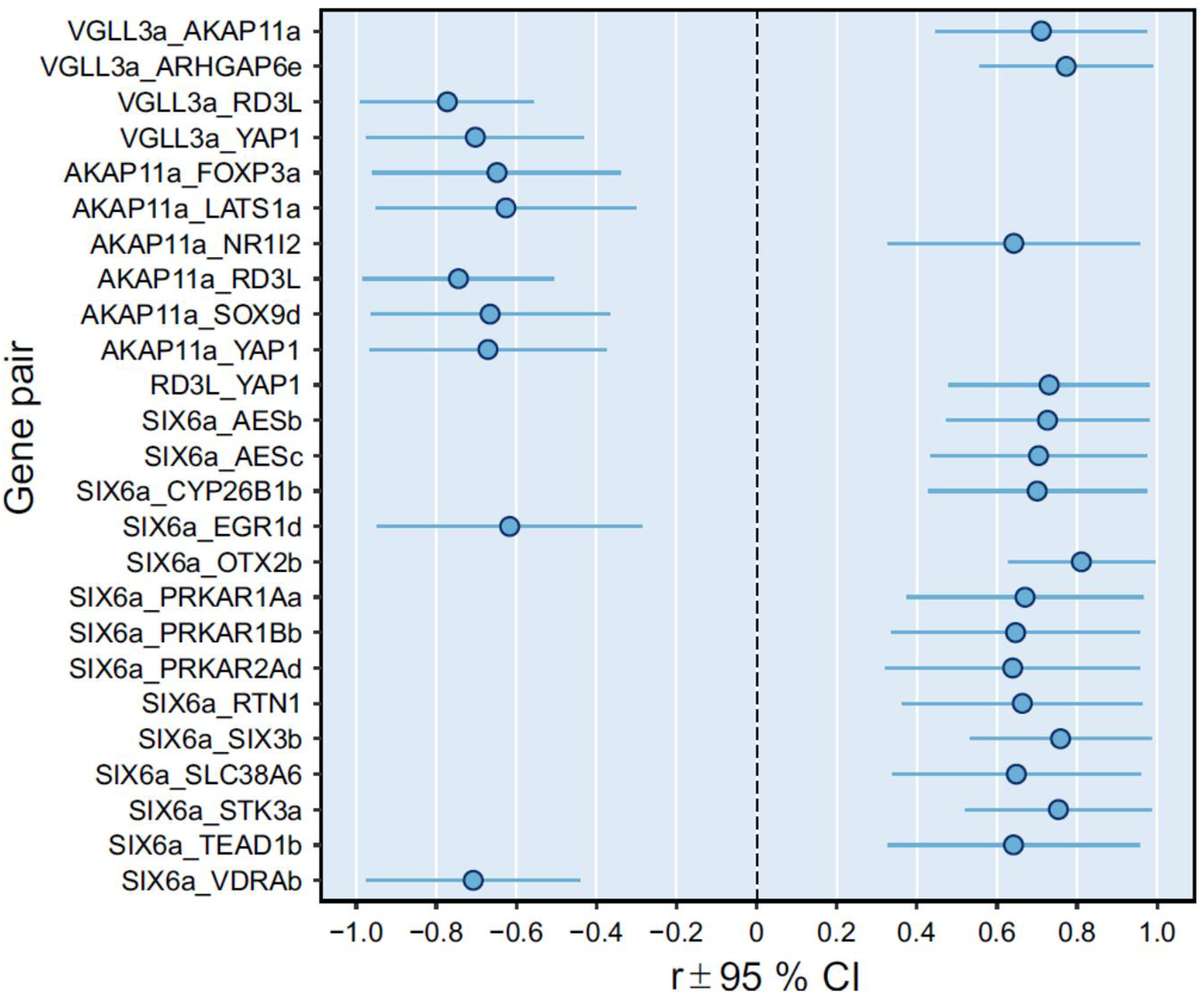
Genes correlating with the age-at-maturity-associated genes during early Atlantic salmon development. The plot shows the 25 gene-pairs for all 24 genes (out of 178 tested) whose expression levels correlated with expression levels of *vgll3a*, *akap11a* or *six6a* with *P* < 0.01. The data are reported as correlation coefficient (r) ± 95 % confidence interval (CI; 2*se).

### mRNA expression level differences between early developmental stages

Age-at-maturity-associated genes *vgll3a*, *akap11a* and *six6a*, and the genes correlating with them were included in the model to estimate transcription expression differences across early developmental stages. Modelling revealed significant gene (*F*_23, 95.2_ = 877.5, *P* < 0.001), stage (*F*_3, 13.4_ = 31.4, *P* < 0.001) and their interaction effects (*F*_69, 140.9_ = 25.8, *P* < 0.001). Controlled for the *FDR*, pairwise post-hoc comparisons indicated that all of the previously identified genes correlated with the age-at-maturity-associated genes, but *nr1i2* and *rtn1*, had significant expression changes during early developmental stages. *Vgll3a* and *rd3l* were upregulated in alevin compared to 258 °d embryo (Fig. 3a). Expression of *vgll3a*, and also *arhgap6e* and *yap1*, was then decreased in fry compared to alevin (Fig. 3a). *Akap11a* was upregulated in fry compared to alevin (Fig. 3b). In contrast, significant expression changes between developmental stages in genes correlating negatively with *akap11a* tended to be downregulations e.g. *foxp3a* in 258 °d embryo, *sox9d* in alevin, and *lats1a*, *sox9d* and *yap1* in fry, compared to the previous developmental stage (Fig. 3b). *Six6a* was downregulated in alevin and fry, compared to the previous stage (Fig. 3c). Similar trend of decreased expression throughout early development was observed with *aesb*, *aesc*, *otx2b*, *prkar2ad*, *six3b*, *slc38a6*, *stk3a* and *tead1b* that correlated positively with *six6a*, but also with *egr1d* (Fig. 3c). In addition, in alevin, *prkar2ad* and *tead1b* expression increased to a higher level than in 186 °d embryo (Fig. 3c). In contrast, *cyp26b1b* was upregulated in 258 °d embryo, *prkar1aa* in alevin and fry, and *prkar1bb* and *vdrab* in fry, compared to the previous stage (Fig. 3d).

**Fig. 3.**
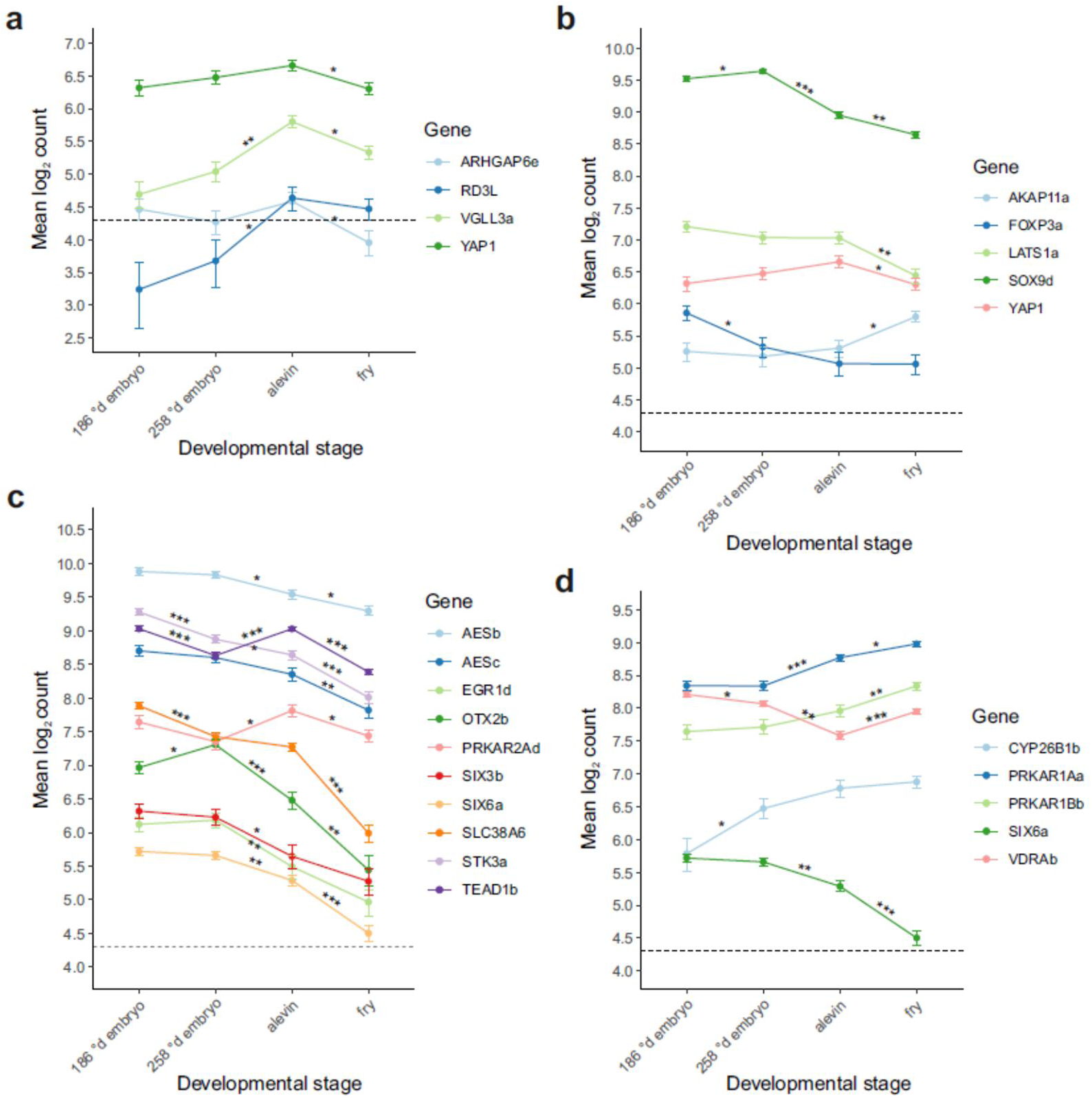
mRNA expression levels of the age-at-maturity-associated genes, and genes correlating with them during early Atlantic salmon development. *Vgll3a* and genes its expression correlates with (**a**), *akap11a* and genes its expression correlates with (**b**) and *six6a* and genes its expression correlates with (**c-d**) in early developmental stages (n = 4). The mixed-model estimates were log_2_-transformed for better visualization and log_2_(20) = 4.3 was defined as a detection threshold. The data are represented as mean ± SE. Expression level differences were tested between all stages but significant comparisons are highlighted by asterisks only for subsequent stages * *FDR*-adjusted *P* < 0.05, ** *P* < 0.01, *** *P* < 0.001

### *Vgll3*, *akap11* and *six6* expression profiles in parr and fry tissues

We characterized mRNA expression patterns of *vgll3, akap11* and *six6* paralogs in 14 different tissues of three-year-old Atlantic salmon parr and blood of fry. *Vgll3a* expression was detected in eight of the 15 tissues: fin (adipose and caudal), gill, heart, liver, muscle, pyloric caeca, spleen and testis (Fig. 4a). Expression of *vgll3b* was only detected in the gill and ovary, the latter having the highest observed tissue-specific expression of *vgll3* paralogs (Fig. 4a). Expression of *akap11a* and *akap11b* was detected in all studied tissues, *akap11b* mostly at a higher level than *akap11a* (Fig. 4c). Interestingly, the highest *akap11a* expression level, higher than that of its paralog, was detected in the ovary (Fig. 4c). *Six6a* expression was observed in four (brain, eye, gill and testis) of the 15 tissues (Fig. 4b), while the *six6b* paralog was expressed only in eye (Fig. 4b).

**Fig. 4.**
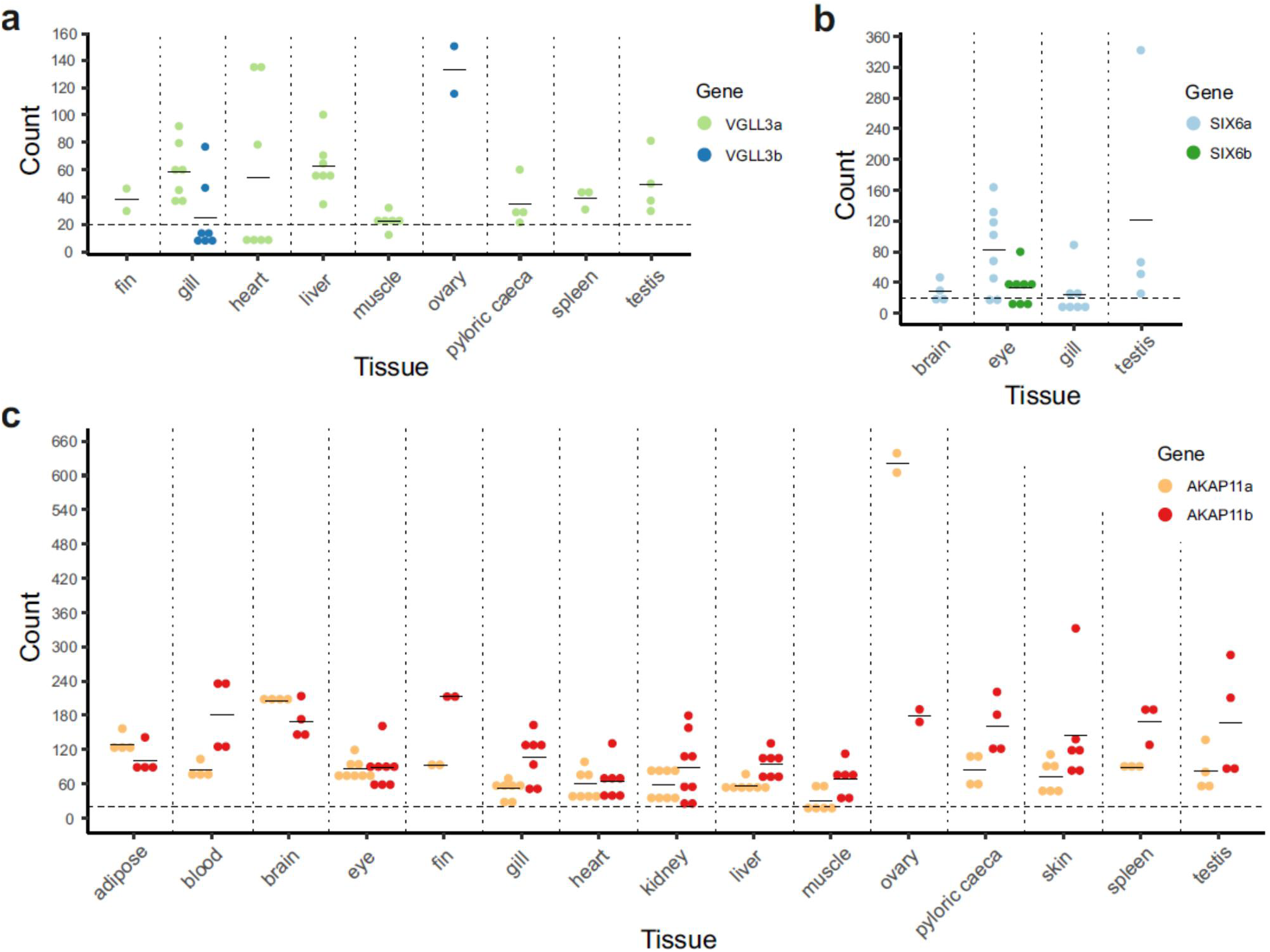
mRNA expression profiles of the age-at-maturity-associated genes and their paralogs in fifteen Atlantic salmon tissues. *Vgll3a* and *vgll3b* (**a**), *six6a* and *six6b* (**b**), and *akap11a* and *akap11b* (**c**) expressed in tissues of three-year old parr and blood of fry. Twenty counts was set as a baseline above which counts are detectable. The data are represented as individual (dot) and mean (line) counts.

## Discussion

We used a custom Nanostring mRNA expression panel designed for Atlantic salmon to examine expression profiles of the *vgll3*, *akap11* and *six6* genes that have been recently associated with sexual maturation timing in Atlantic salmon. Our results show that the age-at-maturity-associated paralogs *vgll3a*, *akap11a* and *six6a*, and also *akap11b*, but not *vgll3b* and *six6b*, are all indeed expressed throughout early embryonic and juvenile salmon development, highlighting the potential relevance of the maturation timing-associated paralogs in developmental processes. We were also able to shed light on which other genes are the potential functional partners of these age-at-maturity-associated genes, and thus provide further insights into the molecular pathways and mechanisms behind maturation. Transcript expression correlations of *vgll3a*, *akap11a* and *six6a* with the subset of 205 prechosen genes related to the functions of the age-at-maturity genes in embryos, alevin and fry suggest that genes associated with salmon maturation timing are linked with cell fate commitment regulation.

Of the studied 205 genes, two (*arhgap6e* and *akap11a*) correlated positively with *vgll3a*, while another two (*yap1* and *rd3l*) correlated negatively. Correlation of *vgll3a* with *arhgap6e* (rho GTPase-activating protein 6) is noteworthy as *ARHGAP6* was shown to be the most downregulated gene in human keratinocytes after *VGLL3* knockdown (54) emphasizing its relevance in Vgll3-induced signaling also in Atlantic salmon. Arhgap6 negatively regulates RhoA (the Rho family GTPase), thus causing actin fiber depolymerization (55). This may be important in the context of maturation-related cellular processes as actin cytoskeleton control is crucial in determining cell fate commitments, such as cell proliferation and differentiation (56). For example, actin cytoskeleton depolymerization is required for cell growth arrest (56) and *sox9* transcriptional activity to induce chondrocyte-specific markers and thus chondrogenesis (57). Accordingly, *vgll3* overexpression upregulates expression of s*ox9* and other genes in chondro- and osteogenesis, and downregulates the main genes in adipogenesis and *vice versa* (20), and, herein, we suggest that Vgll3 could conduct that via actin cytoskeleton regulation. As completion of maturation in Atlantic salmon (58) and another salmon species (59) is highly dependent on the level of fat storage, the expression status of *vgll3* could determine whether to induce adipogenesis or not and, therefore, regulate the timing of maturation. One of the main known regulators of cell fate is the Hippo signalling pathway member Yap (yes associated protein), a transcriptional cofactor that regulates cell proliferation and differentiation based on its cellular location and actin fiber status (56, 60, 61) (Fig. 5). However, there are contradictory results regarding the role of Yap in cell differentiation decisions (62–65). Also, Yap and Vgll3 seem to have overlapping effects on cell fate determination and our suggested outcome of Vgll3 function contradicts with its known role as an inhibitor of adipogenesis (20, 22, 56) (Fig. 5). Nevertheless, the inverse correlation of *vgll3a* with *yap1* in our data implies that these two cofactors could have somewhat opposite roles during development in different stages of cell differentiation processes, such as adipogenesis. And while Yap upregulation has been detected during mammalian puberty (36, 37), it is intriguing that *vgll3*, not *yap1*, genetic variation is tightly associated with maturation timing in Atlantic salmon (6, 66). Moreover, we observed that the expression levels of *vgll3a* and *akap11a* correlate positively, and that expression of *akap11a* further correlates negatively with *yap1*. Interestingly, it is known that protein kinase A (PKA), the regulatory subunit to which Akap11 binds in order to confine the enzyme to discrete locations within the cell (47), can activate the Hippo pathway. It does this by activating Lats kinases, either indirectly by inhibiting Rho GTPase causing actin cytoskeleton depolymerization (67) or directly in response to actin disassembly (68), which leads into inactivation of Yap and further results into inhibition of cell proliferation and induction of adipogenesis. Our results showing inverse correlation between *akap11a* and *lats1a* confirms that PKA-induced Lats regulation is indeed dependent on Akap11. The association of Akap11 with PKA-induced adipogenesis is further supported by our finding that *akap11a* expression correlated negatively with *sox9d*, the downregulation of which is required for adipogenesis (69). The relationship between Vgll3 and PKA-regulating Akap11 appears complex, but the observation that expression of these two age-at-maturity-associated genes correlate with each other and the evidence that they take part in the Hippo pathway speaks strongly for cell fate regulation being an important player in maturation. Fig. 5 outlines a hypothetical model for the regulation of the Hippo pathway by Vgll3 and Akap11 based on a combination of our results and earlier research.

**Fig. 5.**
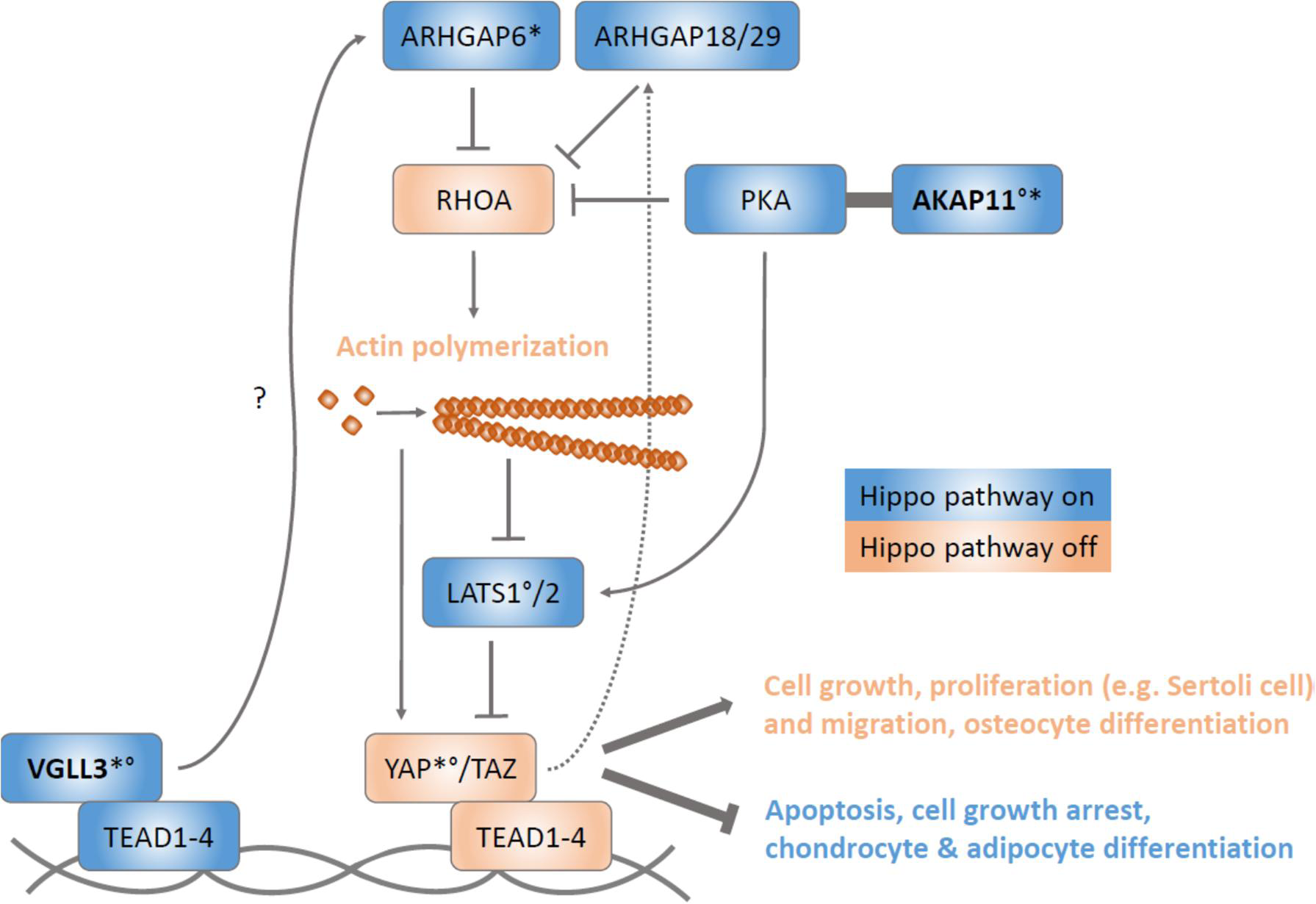
A model for the regulation of the Hippo pathway by Vgll3 and Akap11. By binding to Tead transcription factors, our results suggest that Vgll3 induces expression of Arhgap6 (rho GTPase-activating protein 6) either directly or via other signaling partners. Arhgap6 inhibits RhoA (the Rho family GTPase) activity and thus causes actin fibre depolymerization, which further leads into Lats1/2 activation. Lats kinases phosphorylate the Yap/Taz complex excluding it from the nucleus and causing it to be degraded in the cytoplasm. In addition, PKA (protein kinase A) assisted by Akap11 stimulates Hippo signaling by both inducing and responding to actin depolymerization by activating Lats1/2 kinases either indirectly via RhoA inhibition or directly, respectively. Inactivation of Yap/Taz results in endothelial or epithelial cells to have growth arrest or apoptosis and mesenchymal stem cells (MSCs) to differentiate into chondrocytes or adipocytes. Instead, when the Hippo pathway is inactivated by actin polymerization, Yap/Taz accumulates in the nucleus, binds Teads and induces gene transcription leading to cell growth, proliferation and migration, and osteocyte differentiation. Yap can regulate its own activation via negative feedback loop by upregulating at least Arhgap18 and Arghap29 which suppress RhoA activity. Based on the results of the current study and (56, 61, 64, 67, 68, 70, 71). * Correlation of *vgll3a* mRNA expression with that of *arhgap6e* and *akap11a* (positive), and *yap1* (negative). ° Correlation of *akap11a* mRNA expression with that of *vgll3a* (positive), and *lats1a* and *yap1* (negative).

As *akap11a* is the adjacent gene of *vgll3a* in Atlantic salmon with a distance of 57 kb in between, their transcriptional correlation also indicates that they may be under the same transcriptional regulation and hence may be co-evolving, which could be meaningful when genes share the same molecular pathway (72, 73). Co-expression of adjacent gene pairs is shown to be conserved across eukaryotes (74), also in zebrafish (75). In addition to the occurrence of missense SNPs in both the *vgll3a* and *akap11a* coding regions, the most highly age-at-maturity-associated SNP in the Atlantic salmon *vgll3a* locus resides in a non-coding region between *vgll3a* and *akap11a* (6). This led us to hypothesize that these genes could potentially share a regulatory region between them as it has indeed been shown that a bidirectional promoter can control transcription of an adjacent gene pair (76). Another intriguing link between *vgll3a*, *akap11a* and *yap1* is *rd3l* (retinal degeneration 3-like), which correlated negatively with *vgll3a* and *akap11a* and positively with *yap1*. Knowledge of the functional role(s) of *rd3l* was extremely limited prior to this study. Expression of both *vgll3a* and *rd3l* was significantly upregulated at the alevin stage suggesting that their overall expression is induced simultaneously during development, albeit being oppositely regulated at the individual level. High expression of *vgll3a* in alevin may indicate that Vgll3 starts to induce chondro- and osteogenetic pathways during this stage when lepidotrichia, the bony segmented fin rays, are formed in salmon (50). However, it remains to be studied if and how Rd3l possibly participates in cell fate regulation. Altogether, our data suggest that the functions of *vgll3a* and *akap11a* are linked and associate with cell fate determination by regulating actin cytoskeleton assembly.

Six6 and two other homeobox transcription factor proteins, Six3 and Otx2, regulate the HPG axis signaling in the hypothalamus and pituitary, and eye development (40, 43, 44, 77, 78). In accordance with these studies, our results show high correlation of *six6a* with *six3b* and *otx2b*, and with *aesb* and *aesc*, which encode a corepressor for Six6 and Six3 (79, 80). Further, our results indicate that embryonic stages are crucial time points for the HPG axis and eye development in salmon since *six6a*, *six3b*, *otx2b*, *aesb* and *aesc* were all expressed at their highest level during embryonic stages, after which they started to decline. In addition, we detected that expression of *slc38a6*, which possibly encodes a transporter for glutamine-glutamate cycle in the brain (81) both correlated and covaried temporally with *six6a* during early development. Glutamate is the most prevalent neurotransmitter in the central nervous system, including vertebrate retina (82) and the HPG axis where it induces GnRH release (83). In addition to *slc38a6*, *rtn1*, another gene related to neuroendocrine secretion, correlated with *six6a*. *Rtn1* is highly expressed in the brain, which our results confirmed in salmon parr (data not shown), where it is considered to be a marker for neuronal differentiation (84). Interestingly, both *slc38a6* and *rtn1* are located on the same salmon chromosomal region as *six6a*. Specifically, *slc38a6* locates 123 kb downstream and *rtn1* 165 kb upstream of *six6a*, suggesting that these three genes, which all have roles in neuroendocrine secretion, may be co-regulated, possibly by chromatin modification or folding (73). *Six6a* also correlated and covaried temporally during early development with two Hippo signaling pathway genes, *stk3a* and *tead1b*. These genes encode Mst2 kinase needed to inactivate Yap and a transcription factor partner for Vgll3 and Yap, respectively. Additionally, *six6a* correlated with three PKA regulatory subunit genes. It is known that Hippo signaling regulates eye development (85) and that it is activated in embryonic and postnatal pituitary gland (86).

However, to our knowledge, there are no studies so far linking Six6/Six3 and the Hippo pathway. Instead, it has been shown that Six3, in the embryonic brain (87) and in eye together with Six6 (88), and Hippo signaling (89) repress Wnt signaling, another pathway regulating cell fate (90). Based on our results, it is intriguing that all three Atlantic salmon age-at-maturity-associated genes seem to associate with cell fate determination, especially with the Hippo pathway.

In addition to gaining clues about the early development transcription profiles and molecular mechanisms of *vgll3a, akap11a* and *six6a*, we scrutinized tissue-specific expression patterns of these genes and their paralogs in several tissues of three-year-old Atlantic salmon parr. Our results let us to hypothesize that Vgll3 influences maturation timing by regulating both adipogenesis and gametogenesis. Of the studied samples, *vgll3a* expression was found in fin, gill, muscle, heart, spleen, liver, pyloric caeca – an organ aiding to absorb nutrients specifically in fish – and testis, whereas *vgll3b* expression was restricted to ovary and gill. This, especially sex-dependent gonad-expression pattern with *vgll3a* expressed in testis, but *vgll3b* expressed in ovary, provides evidence for *vgll3* paralog sub-functionalization in salmon. Our results are mostly in line with studies performed in other vertebrates (25, 28, 29) but some differences occur. Knowledge of *vgll3* paralog expression patterns helps to resolve how Vgll3 conducts its function during the maturation process. Based on our results, Vgll3 may participate in inhibiting cell proliferation and organ growth via actin cytoskeleton regulation. This is supported by the study of Kjaerner-Semb et al. (29) who shows that *vgll3a* is expressed in Sertoli cells and downregulated along with some Hippo pathway genes in mature salmon testis, thereby potentially inhibiting the Hippo signaling pathway and inducing subsequent proliferation of Sertoli cells, which are required to provide support for the developing germ cells (91). However, their finding of *vgll3a* being upregulated in granulosa cells of early and late puberty ovary (29) does not support the same mechanism in ovary. This, combined with results of the current study, may indicate that gonad development during maturation is differently regulated in the two sexes. In other tissues, Vgll3 may also function as a regulator for cell fate commitment. According to a previous study (20) and our tissue expression and correlation results, Vgll3 could induce chondro- and osteogenic pathways in fin and gill, which would be needed in growing fish. However, in muscle, where we detected very low *vgll3a* expression, it has been shown that, although Vgll3 induces myogenesis, it is expressed at low levels in healthy muscle (22). In line with this, our finding that adipose tissue lacks both *vgll3* paralog expression is reasonable, as it is known that knockdown of *vgll3* expression promotes pre-adipocytes to differentiate into mature adipocytes (20). In other words, also in salmon, decreased expression of *vgll3a* may be required to induce adipogenesis, which is critical to ensure enough fat-derived energy for sexual maturation (58, 59).

Unlike the *vgll3* genes, both *akap11* paralogs were expressed in all studied tissues, which is consistent with Reinton et al. (47), emphasizing the relevance of Akap11 in basic cellular functions. However, *akap11b* was expressed at a higher level than *akap11a* in most tissues, suggesting its higher functional activity. Our results in salmon gonads confirm the ubiquitous nature of *akap11* expression but further suggests somewhat specialized functions for each paralog. In contrast to *vgll3* whose age-at-maturity-associated paralog was expressed in testis and other paralog in ovary, both *akap11* paralogs were expressed in both ovary and testis. However, the age-at-maturity-associated *akap11a* and several paralogs encoding the regulatory subunits of PKA were more highly expressed in ovary, whereas *akap11b* expression was higher in testis. This differs somewhat from the study by Reinton, Collas (47) in that *akap11* mRNA expression in the human ovary was extremely low. So far, it is known that as important as in sustaining interphase during mitosis (48), PKA activation is required to maintain meiotic arrest of oocyte (92). Since the studied salmon ovaries were immature, high *akap11a* and PKA regulatory subunit mRNA expression could result from the need of PKA to sustain oocytes in meiotic arrest. In testis, however, the Akap11/PKA complex is suggested to have a dual role in meiosis during spermatogenesis and in motility of mature sperm (47). Overall, our results propose that, in Atlantic salmon, *akap11* has evolved paralog-specific functional differences, especially in gonads.

Expression of *six6a* was detected in brain, eye, testis and gill. To our knowledge, expression in gill has not been reported earlier. Instead, expression of *six6b* was detected only in eye, which could imply that the expression patterns observed for the age-at-maturity-associated paralog have evolved and expanded to the HPG axis-related tissues and, thus, reproduction. Although expression of *SIX6* has been earlier detected in testis (42), its specific function remains unknown. Another transcript expressed in brain, but whose expression we detected also in testis, is *cgba* (data not shown). This gene encodes the beta-subunit of Fsh (follicle-stimulating hormone) gonadotropin, which is normally expressed in pituitary and regulated by Six6 (43). These findings hint that Six6 may potentially regulate the expression of the testicular Fsh beta subunit and thus testis development. This is supported by an earlier observation that Fsh beta subunit mRNA is expressed in mouse testis, where it is suggested to play paracrine or autocrine role in the regulation of testicular function (93), such as Sertoli and germ cell development (94).

We conducted one of the rare follow-up studies of a GWAS association in a wild animal species with a view to better understanding the molecular mechanisms of the previously discovered genotype-phenotype association. Our temporal assessment of mRNA expression of Atlantic salmon age-at-maturity-associated *vgll3a*, *akap11a* and *six6a* during five developmental stages revealed differently regulated expression in a development stage- and tissue-specific manner. Co-expression analysis of the age-at-maturity-associated genes and 205 other pre-selected genes indicated co-regulation of *vgll3a* and *akap11a*, and a novel role for Vgll3 in regulating actin cytoskeleton assembly – a process required in cell proliferation and differentiation – and that this regulation could be assisted by Akap11-directed PKA function. In addition, we were able to confirm the same expressional pattern in salmon that *six6* and its partners have in the HPG axis and eye development regulation in other vertebrates. Moreover, we propose that Six6 may associate more broadly with neuroendrocrine regulation in the HPG axis and have a direct role in testis function. Further, our data provide the first evidence that both Vgll3 and Six6 paralogs have sub-functionalized roles in different tissues. Overall, we conclude that Vgll3, Akap11 and Six6 may contribute to influencing Atlantic salmon maturation timing via affecting on adipogenesis and gametogenesis by regulating cell fate commitment and the HPG axis. Further studies are required to determine the more specific cellular molecular mechanisms of Vgll3, Akap11 and Six6 in sexual maturation processes. This work provides important information for guiding such work, also in other organisms.

## Abbreviations

Fsh: Follicle-stimulating hormone
GnRH: Gonadotropin-releasing hormone
GWAS: Genome-wide association study
HPG: Hypothalamic-pituitary-gonadal
PKA: protein kinase A
RhoA: Rho family GTPase
SNP: Single nucleotide polymorphism

## Ethics approval and consent to participate

Salmon were reared under animal experimentation licenses obtained by the Natural Resources Institute Finland and University of Helsinki.

## Authors’ contributions

JK, PVD and CRP designed the study. JE provided access to the animals. JK, PVD, AH, TA, JE and CRP collected the samples. JK conducted the experiments. JK and PVD analyzed the data. JK interpreted the analyzed data. JK and CRP wrote the first draft of the manuscript. All authors have revised and approved the submitted version of the manuscript.

## Acknowledgements

We kindly thank Päivi Anttonen and Risto Kannel for help collecting the samples and providing rearing temperature data. We also acknowledge Katja Salminen for help extracting RNA and Victoria Pritchard for suggesting genes for the mRNA expression panel.

