## Additional file 1 for "Transcription profiles of age-at-maturity-associated genes suggest cell fate commitment regulation as a key factor in the Atlantic salmon maturation process"

Additional table 1. Tissues of eight fish chosen for the study.

| Fish ID | Sex | Habitat | Maturity stage | Tissue | | | | | | | | | | | | | | |
|---|---|---|---|---|---|---|---|---|---|---|---|---|---|---|---|---|---|---|
| | | | | Adipose | Brain | Eye | Fin (adipose) | Fin (tail) | Gill | Heart | Kidney | Liver | Muscle | Ovary | Pyloric caeca | Skin | Spleen | Testis |
| N.1 | Male | Hatchery | Immature | x | x | x | x | x | x | x | x | x | x | | x | x | x | |
| N.2 | Male | Hatchery | Immature | x | x | x | | | x | x | x | x | | | x | | | |
| N.3 | Female | Hatchery | Immature | x | x | x | | | x | x | x | x | x | x | x | x | x | x |
| N.4 | Female | Hatchery | Immature | x | x | x | | | x | | x | x | x* | x | x | x | x | x |
| I.1 | Male | Wild | Immature | | | x | | | x | x | x | x* | x | | | | | |
| I.2 | Male | Wild | Immature | | | x | | | x | x | x | x | x | | | x | | |
| I.3 | Male | Wild | Mature | | | x | | | x | x | x | x | x | | | x | | x |
| I.4 | Male | Wild | Mature | | | x | | | | x | x | x | x | | | x | | x |

N = Neva, I = Inarijoki

* Samples were excluded from the data due to not passing the quality control during data analysis
