## Additional file 2 for "Transcription profiles of age-at-maturity-associated genes suggest cell fate commitment regulation as a key factor in the Atlantic salmon maturation process"

Kurko et al. Additional file 2. Genes included in Nanostring panel

| Accesion number | Gene ID | Gene name | Functional category |
|---|---|---|---|
| XR_001323797.1 | ADMa | ADM-like/Adrenomedullin-like | HPG axis, Cell fate |
| XM_014175339.1 | ADMb | ADM-like/Adrenomedullin-like | HPG axis, Cell fate |
| XM_014126713.1 | ADMc | ADM-like/Adrenomedullin-like | HPG axis, Cell fate |
| XM_014161791.1 | AEBP1 | AE (Adipocyte enhancer) binding protein 1 | Metabolism, Cell fate |
| XM_014150407.1 | AESa | Amino-terminal enhancer of split-like | HPG axis |
| XM_014216422.1 | AESb | Amino-terminal enhancer of split-like | HPG axis |
| XM_014169692.1 | AESc | Amino-terminal enhancer of split-like | HPG axis |
| XM_014173902.1 | AKAP11a | A-Kinase (PRKA) anchor protein 11 | Cell fate, Chr 25 candidate region |
| XM_014164979.1 | AKAP11b | A-Kinase anchor Protein 11-like | Cell fate, Chr 25 candidate region |
| XM_014201564.1 | APOC1a | Apolipoprotein C-I | Metabolism |
| XM_014155604.1 | APOC1b | Apolipoprotein C-I-like | Metabolism |
| XM_014137515.1 | ARB1 | Androgen receptor-like | HPG axis |
| XM_014196828.1 | ARB2/AR | Androgen receptor | HPG axis |
| XM_014150285.1 | ARHGAP6a | rho GTPase-activating protein 6-like | Cell fate |
| XM_014164375.1 | ARHGAP6b | rho GTPase-activating protein 6-like | Cell fate |
| XM_014151712.1 | ARHGAP6c | rho GTPase-activating protein 6-like | Cell fate |
| XM_014156305.1 | ARHGAP6d | rho GTPase-activating protein 6-like | Cell fate |
| XM_014174484.1 | ARHGAP6e | rho GTPase-activating protein 6-like | Cell fate |
| XM_014207436.1 | ARHGAP6f | rho GTPase-activating protein 6-like | Cell fate |
| XM_014173895.1 | ASMT | Acetylserotonin O-methyltransferase | Metabolism, Cell fate, Chr 25 candidate region |
| XM_014198381.1 | CADM2A | Cell adhesion molecule 2-like | Metabolism |
| XM_014129854.1 | CADM2Ba | Cell adhesion molecule 2-like | Metabolism |
| XM_014163653.1 | CADM2Bb | Cell adhesion molecule 2-like | Metabolism |
| XM_014173899.1 | CCDC138 | Coiled-coil domain-containing protein 138-like | Chr 25 candidate region |
| NM_001139931.1 | CEBPAa | CCAAT/enhancer binding protein (C/EBP), alpha | Cell fate |
| XM_014127824.1 | CEBPAb | CCAAT/enhancer-binding protein alpha-like | Cell fate |
| NM_001139913.1 | CEBPBa | CCAAT/enhancer binding protein (C/EBP), beta | Cell fate |
| XM_014146703.1 | CEBPBb | CCAAT/enhancer-binding protein beta-like | Cell fate |
| NM_001141493.2 | CEBPDa | CCAAT/enhancer binding protein (C/EBP), delta | Cell fate |
| XM_014191131.1 | CEBPDb | CCAAT/enhancer binding protein (C/EBP), delta | Cell fate |
| XM_014210156.1 | CGAa | Glycoprotein hormones alpha chain 1-like | HPG axis |

| | | | |
|---|---|---|---|
| XM_014209152.1 | CGAb | Glycoprotein hormones alpha chain 2-like | HPG axis |
| XM_014126340.1 | CGBa | Gonadotropin subunit beta-1 | HPG axis |
| XM_014179923.1 | CGBb/GTHB2 | Gonadotropin subunit beta-2 | HPG axis |
| XM_014173900.1 | CHMP2B | Charged multivesicular body protein 2b-like | Chr 25 candidate region |
| XM_014204893.1 | COL12A1a | Collagen, type XII, alpha 1 | Cell fate |
| XM_014144106.1 | COL12A1b | Collagen alpha-1(XII) chain-like | Cell fate |
| XM_014210332.1 | COL12A1c | Collagen alpha-1(XII) chain-like | Cell fate |
| XM_014204457.1 | COL1A1a | Collagen alpha-1(I) chain | Cell fate |
| XM_014192569.1 | COL1A1b | Collagen alpha-1(I) chain-like | Cell fate |
| XM_014178512.1 | COL1A2 | Collagen, type I, alpha 2 | Cell fate |
| XM_014209621.1 | CXXC4 | CXXC finger protein 4 | Cell fate |
| XM_014156772.1 | CYP26B1a | Cytochrome P450 26B1 | Cell fate |
| XM_014206393.1 | CYP26B1b | Cytochrome P450 26B1-like | Cell fate |
| XM_014210736.1 | DHRS7 | Dehydrogenase/reductase (SDR family) member 7 | Metabolism, Chr 9 candidate region |
| XM_014212308.1 | DLK1a | Delta-like 1 homolog (Drosophila)/PREF1 | Cell fate |
| XM_014154083.1 | DLK1b | Delta-like 1 homolog (Drosophila)/PREF2 | Cell fate |
| XM_014173644.1 | DNAJC15 | DnaJ (Hsp40) homolog, subfamily C, member 15 | Chr 25 candidate region |
| XM_014173897.1 | EDAR | Tumor necrosis factor receptor superfamily member EDAR-like/Ectodysplasin A Receptor | Chr 25 candidate region |
| NM_001141824.1 | EGR1a | Early growth response 1 | HPG axis, Cell fate, Metabolism |
| XM_014198129.1 | EGR1b | Early growth response protein 1-like | HPG axis, Cell fate, Metabolism |
| XM_014212603.1 | EGR1c | Early growth response protein 1-like | HPG axis, Cell fate, Metabolism |
| XM_014200066.1 | EGR1d | Early growth response protein 1-like | HPG axis, Cell fate, Metabolism |
| XM_014173914.1 | ENOX1 | Ecto-NOX disulfide-thiol exchanger 1 | Cell fate, Chr 25 candidate region |
| XM_014173912.1 | EPSTI1 | Epithelial stromal interaction 1 (breast) | Chr 25 candidate region |
| XM_014205563.1 | ESR1 | Estrogen receptor 1 | HPG axis, Cell fate |
| XM_014196428.1 | ESRRAa | Steroid hormone receptor ERR1-like/Estrogen related receptor alpha | Metabolism, Cell fate |
| XM_014129095.1 | ESRRAb | Steroid hormone receptor ERR1-like/Estrogen related receptor alpha | Metabolism, Cell fate |
| XM_014210759.1 | FGFRL1 | Fibroblast growth factor receptor-like 1 | Cell fate, Chr 9 candidate region |
| XM_014146077.1 | FHAD1 | Forkhead-associated domain-containing protein 1-like | Cell fate |
| XM_014195874.1 | FOXO1a | Forkhead box protein O1-A-like | Cell fate |
| XM_014123816.1 | FOXO1b | Forkhead box protein O1-A-like | Cell fate |
| XM_014163663.1 | FOXO1c | Forkhead box protein O1-A-like | Cell fate |
| NM_001198847.1 | FOXP3a | Forkhead box P3 | HPG axis |

| | | | |
|---|---|---|---|
| XM_014146362.1 | FOXP3b | Forkhead box protein P1-B-like | HPG axis |
| NM_001126230.1 | FSHR | Follicle stimulating hormone receptor | HPG axis |
| XM_014148799.1 | FSTL3a | Follistatin-related protein 3-like | Cell fate |
| XM_014216321.1 | FSTL3b | Follistatin-related protein 3-like | Cell fate |
| XM_014216042.1 | GADD45Ba | Growth arrest and DNA damage-inducible protein GADD45 beta-like | Cell fate, Metabolism |
| XM_014149394.1 | GADD45Bb | Growth arrest and DNA damage-inducible protein GADD45 beta-like | Cell fate, Metabolism |
| NM_001123676.1 | GHa | Growth hormone prepeptide | Metabolism, HPG axis |
| XM_014204437.1 | GHb | Somatotropin-2/Growth hormone | Metabolism, HPG axis |
| XM_014133894.1 | GHR1 | Growth hormone receptor isoform 1 precursor | Metabolism, HPG axis |
| NM_001139585.1 | GHRL | Ghrelin and obestatin prepropeptide | Metabolism, HPG axis |
| XM_014150371.1 | GHSRa | Growth hormone secretagogue receptor 1 | Metabolism, HPG axis |
| XM_014151925.1 | GHSRb | Growth hormone secretagogue receptor type 1-like | Metabolism, HPG axis |
| XM_014202065.1 | GHSRc | Growth hormone secretagogue receptor type 1-like | Metabolism, HPG axis |
| NM_001146456.1 | GLHA1 | Glycoprotein hormones alpha chain 1 | HPG axis |
| XM_014195502.1 | GLHA2a | Pituitary alpha-2 glycoprotein hormone subunit precursor | HPG axis |
| XM_014209153.1 | GLHA2b | Glycoprotein hormones alpha chain 2-like | HPG axis |
| XM_014154458.1 | GNRH | Gonadotrophin releasing hormone | HPG axis |
| XM_014160183.1 | GNRH1 | Gonadotropin-releasing hormone 1 (luteinizing-releasing hormone) | HPG axis |
| XM_014142277.1 | GNRHR1a | Gonadotropin-releasing hormone II receptor-like/Gonadotropin-releasing hormone 1 receptor | HPG axis |
| XM_014177690.1 | GNRHR1b | Gonadotropin-releasing hormone receptor 1 | HPG axis |
| XM_014148656.1 | GNRHR4a | Gonadotropin-releasing hormone II receptor-like/Gonadotropin releasing hormone receptor 4 | HPG axis |
| XM_014124885.1 | GNRHR4b | Gonadotropin-releasing hormone II receptor-like/Gonadotropin releasing hormone receptor 4 | HPG axis |
| XM_014161730.1 | GNRHRa | Gonadotropin-releasing hormone II receptor-like | HPG axis |
| XM_014201132.1 | GNRHRb | Gonadotropin-releasing hormone II receptor-like | HPG axis |
| XM_014210245.1 | GREB1La | Protein GREB1-like/Growth regulation by estrogen in breast cancer 1 like/GREB1 like retinoic acid receptor coactivator | HPG axis |
| XM_014142369.1 | GREB1Lb | Protein GREB1-like/Growth regulation by estrogen in breast cancer 1 like/GREB1 like retinoic acid receptor coactivator | HPG axis |
| XM_014210723.1 | HIF1A | Hypoxia-inducible factor 1-alpha-like | Cell fate, Chr 9 candidate region |
| XM_014173643.1 | HTR1F | 5-hydroxytryptamine receptor 1F-like | Chr 25 candidate region |
| XM_014167209.1 | KDM5Ba | Lysine-specific demethylase 5B-B-like | Cell fate |
| XM_014132540.1 | KDM5Bb | Lysine-specific demethylase 5B-B-like | Cell fate |
| XM_014136221.1 | KDM5Bc | Lysine-specific demethylase 5B-B-like | Cell fate |
| XM_014146695.1 | KDM5Bd | Lysine-specific demethylase 5B-B-like | Cell fate |
| XM_014210754.1 | KIF15 | Kinesin family member 15 | Chr 9 candidate region |

| | | | |
|---|---|---|---|
| XM_014132950.1 | KLF15a | Krueppel-like factor 15 | Cell fate |
| XM_014132138.1 | KLF15b | Krueppel-like factor 15 | Cell fate |
| XM_014168045.1 | KLF15c | Krueppel-like factor 15 | Cell fate |
| XM_014135694.1 | KLF15d | Krueppel-like factor 15 | Cell fate |
| XM_014143948.1 | LAPTM4Aa | Lysosomal protein transmembrane 4 alpha | Cell fate |
| NM_001140354.1 | LAPTM4Ab/LAP4A | Lysosomal-associated transmembrane protein 4A | Cell fate |
| XM_014205037.1 | LATS1a | Serine/threonine-protein kinase LATS1-like | Cell fate |
| XM_014144217.1 | LATS1b | Serine/threonine-protein kinase LATS1-like | Cell fate |
| XM_014164783.1 | LATS2a | Serine/threonine-protein kinase LATS2-like | Cell fate |
| XM_014174046.1 | LATS2b | Serine/threonine-protein kinase LATS2-like | Cell fate |
| NM_001279134.1 | LEPB1 | Leptin B1 | Metabolism, HPG axis |
| XM_014123299.1 | LEPRa | Leptin receptor | Metabolism, HPG axis |
| XM_014168840.1 | LEPRb | Leptin receptor-like | Metabolism, HPG axis |
| XM_014179976.1 | LHR/LHCGR | Luteinizing hormone receptor | HPG axis |
| XM_014210738.1 | LRRC9 | Leucine-rich repeat-containing protein 9-like | Chr 9 candidate region |
| XM_014140480.1 | MC4Ra | Melanocortin receptor 4-like | Metabolism |
| XM_014190362.1 | MC4Rb | Melanocortin receptor 4-like | Metabolism |
| XM_014180569.1 | MC4Rc | Melanocortin receptor 4-like | Metabolism |
| XM_014157590.1 | MC4Rd | Melanocortin receptor 4-like | Metabolism |
| NM_001140524.1 | MMP13a | Collagenase 3/Matrix metallopeptidase 13 | Cell fate |
| XM_014137820.1 | MMP13b | Collagenase 3-like /Matrix metallopeptidase 13 | Cell fate |
| XM_014137819.1 | MMP13c | Collagenase 3-like /Matrix metallopeptidase 13 | Cell fate |
| XM_014210730.1 | MNAT1 | MNAT CDK-activating kinase assembly factor 1 | Cell fate, Chr 9 candidate region |
| XM_014133756.1 | MST1a | Hepatocyte growth factor-like protein/ Macrophage stimulating 1 | Cell fate |
| XM_014166189.1 | MST1b | Hepatocyte growth factor-like protein /Macrophage stimulating 1 | Cell fate |
| NM_001123644.1 | MYF5a | Myogenic factor 5 | Cell fate |
| XM_014153440.1 | MYF5b | Myogenic factor 5-like | Cell fate |
| XM_014174162.1 | NR1I2/PXR | Nuclear receptor subfamily 1 group I member 2-like | Metabolism |
| XM_014210818.1 | OTX2a | Orthodenticle homeobox 2 | HPG axis |
| XM_014185285.1 | OTX2b | Homeobox protein OTX2 | HPG axis |
| XM_014144803.1 | OTX2c | Homeobox protein OTX2-like | HPG axis |
| XM_014205787.1 | OTX2d | Homeobox protein OTX2-like | HPG axis |
| XM_014210604.1 | PCNXL4 | Pecanex-like 4 (Drosophila) | Chr 9 candidate region |

| | | | |
|---|---|---|---|
| NM_001252363.1 | PGRa | Nuclear progesterone receptor | HPG axis |
| XM_014198242.1 | PITX1a | Paired-like homeodomain 1 | HPG axis, Cell fate |
| XM_014123791.1 | PITX1b | Paired-like homeodomain transcription factor 1beta | HPG axis, Cell fate |
| XM_014124069.1 | PPARAa | Peroxisome proliferator-activated receptor alpha | Metabolism |
| XM_014169858.1 | PPARAb | Peroxisome proliferator-activated receptor alpha-like | Metabolism |
| XM_014152661.1 | PPARAc | Peroxisome proliferator-activated receptor alpha-like | Metabolism |
| XM_014167444.1 | PPARB1A | Peroxisomal proliferator-activated receptor beta1A | Metabolism |
| XM_014147251.1 | PPARB2B | Peroxisome proliferator-activated receptor beta2B | Metabolism |
| NM_001123546.1 | PPARGa | Peroxisome proliferator activated receptor gamma | Metabolism |
| XM_014135113.1 | PPARGb | Peroxisome proliferator-activated receptor gamma-like | Cell fate, Metabolism |
| XM_014168482.1 | PPARGc | Peroxisome proliferator-activated receptor gamma-like | Cell fate, Metabolism |
| XM_014210734.1 | PPM1A | Protein phosphatase 1A-like | Cell fate, Chr 9 candidate region |
| XM_014179785.1 | PRKAR1Aa | Protein kinase, cAMP-dependent, regulatory, type I, alpha | Cell fate, Metabolism, Housekeeping |
| XM_014209627.1 | PRKAR1Ab | cAMP-dependent protein kinase type I-alpha regulatory subunit | Cell fate, Metabolism |
| XM_014202328.1 | PRKAR1Ac | cAMP-dependent protein kinase type I-alpha regulatory subunit-like | Cell fate, Metabolism |
| XM_014202670.1 | PRKAR1Ba | cAMP-dependent protein kinase type I-beta regulatory subunit-like | Cell fate, Metabolism |
| XM_014202711.1 | PRKAR1Bb | cAMP-dependent protein kinase type I-beta regulatory subunit-like | Cell fate, Metabolism |
| XM_014131913.1 | PRKAR2Aa | cAMP-dependent protein kinase type II-alpha regulatory subunit-like | Cell fate |
| XM_014167675.1 | PRKAR2Ab | cAMP-dependent protein kinase type II-alpha regulatory subunit-like | Cell fate |
| XM_014135164.1 | PRKAR2Ac | cAMP-dependent protein kinase type II-alpha regulatory subunit-like | Cell fate |
| XM_014146325.1 | PRKAR2Ad | cAMP-dependent protein kinase type II-alpha regulatory subunit-like | Cell fate |
| XM_014125918.1 | PRKAR2Ba | protein kinase, cAMP-dependent, regulatory, type II, beta | Cell fate, Metabolism |
| XM_014147660.1 | PRKAR2Bb | cAMP-dependent protein kinase type II-beta regulatory subunit-like | Cell fate, Metabolism |
| XM_014152968.1 | PRKAR2Bc | cAMP-dependent protein kinase type II-beta regulatory subunit-like | Cell fate, Metabolism |
| XM_014207731.1 | PRKAR2Bd | cAMP-dependent protein kinase type II-beta regulatory subunit-like | Cell fate, Metabolism |
| XM_014210726.1 | PRKCH | Protein kinase C eta type | Chr 9 candidate region |
| XM_014210752.1 | RD3L | Protein RD3-like | Chr 9 candidate region |
| XM_014125479.1 | RLN3Aa | Relaxin-3-like | HPG axis, Metabolism |
| XM_014202846.1 | RLN3Ab | Relaxin-3-like | HPG axis, Metabolism |
| XM_014202852.1 | RLN3Ac | Relaxin-3-like | HPG axis, Metabolism |
| XM_014138792.1 | RLN3Ad | Relaxin-3-like | HPG axis, Metabolism |
| XM_014210748.1 | RTN1 | Reticulon 1 | HPG axis, Cell fate, Chr 9 candidate region |
| XM_014172379.1 | RXRAa | Retinoid X receptor, alpha | Metabolism, Cell fate |

| | | | |
|---|---|---|---|
| XM_014128611.1 | RXRAb | Retinoic acid receptor RXR-alpha-A | Metabolism, Cell fate |
| XM_014132501.1 | RXRAc | Retinoic acid receptor RXR-alpha-A-like | Metabolism, Cell fate |
| XM_014141731.1 | RXRBAa | Retinoic acid receptor RXR-beta-A-like | Metabolism, Cell fate, Housekeeping |
| XM_014202184.1 | RXRBAb | Retinoic acid receptor RXR-beta-A-like | Metabolism, Cell fate |
| XM_014177327.1 | RXRBAc | Retinoic acid receptor RXR-beta-A | Metabolism, Cell fate |
| XM_014190174.1 | RXRGAa | Retinoic acid receptor RXR-gamma-A-like | Metabolism, Cell fate |
| XM_014139718.1 | RXRGAb | Retinoic acid receptor RXR-gamma-A-like | Metabolism, Cell fate |
| NM_001139628.1 | SIX1 | Sine oculis homeobox like 1 | Cell fate, Chr 9 candidate region |
| XM_014179983.1 | SIX3a | SIX homeobox 3 | HPG axis |
| XM_014210714.1 | SIX3b | Homeobox protein SIX3-like | HPG axis |
| XM_014158209.1 | SIX3c | Homeobox protein SIX3-like | HPG axis |
| XM_014210729.1 | SIX4 | Homeobox protein SIX4-like | Cell fate, Chr 9 candidate region |
| XM_014210731.1 | SIX6a | SIX homeobox 6 | HPG axis, Chr 9 candidate region |
| XM_014189802.1 | SIX6b | Homeobox protein SIX6 | HPG axis |
| XM_014210728.1 | SLC38A6 | Solute carrier family 38, member 6 | HPG axis, Chr 9 candidate region |
| XM_014134685.1 | SNAI1 | Protein snail homolog Sna-like/Snail family transcriptional repressor 1 | Cell fate |
| NM_001140203.1 | SNAI2a | Snail family transcriptional repressor 2 | Cell fate |
| XM_014140125.1 | SNAI2b | Zinc finger protein SNAI2-like | Cell fate |
| XM_014202610.1 | SOX9a | Transcription factor SOX-9-A-like | Cell fate, HGP axis |
| XM_014194273.1 | SOX9b | Transcription factor SOX-9/Transcription factor SOX-8-like | Cell fate, HGP axis |
| XM_014179186.1 | SOX9c | Transcription factor SOX-9-A-like | Cell fate, HGP axis |
| XM_014158399.1 | SOX9d | Transcription factor SOX-9-B-like | Cell fate, HGP axis |
| NM_001123551.1 | SPP2 | Secreted phosphoprotein 2 | Cell fate |
| XM_014143284.1 | STK3a | Serine/threonine-protein kinase 3-like | Cell fate |
| XM_014178825.1 | STK3b | Serine/threonine-protein kinase 3-like | Cell fate |
| XM_014210746.1 | TDRD9 | Putative ATP-dependent RNA helicase TDRD9 | Cell fate, Chr 9 candidate region |
| XM_014126883.1 | TEAD1a | Transcriptional enhancer factor TEF-1-like | Cell fate |
| XM_014175316.1 | TEAD1b | Transcriptional enhancer factor TEF-1 | Cell fate |
| XM_014125445.1 | TEAD1c | Transcriptional enhancer factor TEF-1-like | Cell fate |
| XM_014148054.1 | TEAD1d | Transcriptional enhancer factor TEF-1-like | Cell fate |
| XM_014126867.1 | TEAD2 | TEA domain family member 2 | Cell fate |
| XM_014133831.1 | TEAD3a | TEA domain family member 3 | Cell fate |
| XM_014166078.1 | TEAD3b | Transcriptional enhancer factor TEF-5-like | Cell fate |

| | | | |
|---|---|---|---|
| XM_014160992.1 | TLE1a | Transducin-like enhancer of split 1 (E(sp1) homolog, Drosophila) | HPG axis |
| XM_014169695.1 | TLE1b | Transducin-like enhancer protein 1 | HPG axis |
| XM_014172558.1 | TLE4a | Transducin-like enhancer of split 4 | HPG axis |
| XM_014216416.1 | TLE4b | Transducin-like enhancer protein 1/Transducin-like enhancer protein 4 | HPG axis |
| XM_014172750.1 | TLE4c | Transducin-like enhancer protein 4 | HPG axis |
| XM_014173909.1 | TNFSF11 | Tumor necrosis factor (ligand) superfamily, member 11 | Cell fate, Chr 25 candidate region |
| XM_014132120.1 | VDRAa | Vitamin D3 receptor B | Metabolism, Cell fate, Housekeeping |
| XM_014146371.1 | VDRAb/VDR0 | Vitamin D receptor | Metabolism, Cell fate, Housekeeping |
| XM_014135458.1 | VDRAc | Vitamin D3 receptor A-like | Metabolism, Cell fate |
| XM_014167996.1 | VDRAd | Vitamin D3 receptor B | Metabolism, Cell fate |
| XM_014173901.1 | VGLL3a | Transcription cofactor vestigial-like protein 3 | Cell fate, Chr 25 candidate region |
| XM_014164976.1 | VGLL3b | Transcription cofactor vestigial-like protein 3 | Cell fate |
| XM_014123631.1 | WWTR1a | WW domain-containing transcription regulator protein 1-like | Cell fate |
| XM_014168664.1 | WWTR1b | WW domain-containing transcription regulator protein 1-like | Cell fate |
| XM_014215593.1 | YAP1 | Transcriptional coactivator YAP1-like | Cell fate |
| XM_014177562.1 | EF1AC | Elongation factor 1-alpha-like | Housekeeping |
| NM_001140843.1 | RPS20 | 40S ribosomal protein S20 (rs20) | Housekeeping |
