## Additional file 3 for "Transcription profiles of age-at-maturity-associated genes suggest cell fate commitment regulation as a key factor in the Atlantic salmon maturation process"

```
#### please define path

SubmissionPath = ""

########################################################
####
####  calculate pairwise correlations
####
#######################################################

# load data
RNADataFrame = NULL
RNADataFrame =
read.table(paste0(SubmissionPath,"NormCountDevelopmentalData.txt"), header=TRUE)
head(RNADataFrame)
dim(RNADataFrame)

# remove transcripts that show below BaselineLimit across all stages
temp = NULL
temp = table(RNADataFrame$Transcript.ID[RNADataFrame$BaselineLimit ==
"detected"])
temp
non.detected.genes = names(temp[temp == 0])
length(unique(non.detected.genes))

dim(RNADataFrame)
# [1] 3816    7
RNADataFrame.detected = RNADataFrame[which(!RNADataFrame$Transcript.ID %in%
non.detected.genes),]
dim(RNADataFrame.detected)
# [1] 3060    7

# make table with present genes
RNADataFrame.detected$Transcript.ID =
as.character(RNADataFrame.detected$Transcript.ID)
selection = unique(RNADataFrame.detected$Transcript.ID)
length(selection)

tempDat = NULL
tempDat = cbind(
        "Sample.ID" =
RNADataFrame.detected$Sample.ID[which(RNADataFrame.detected$Transcript.ID ==
unique(RNADataFrame.detected$Transcript.ID)[1])],
        "Stage"         =
RNADataFrame.detected$Stage[which(RNADataFrame.detected$Transcript.ID ==
unique(RNADataFrame.detected$Transcript.ID)[1])],
        unstack(RNADataFrame.detected, values ~ Transcript.ID))
head(tempDat)

# calculate cors, determine significance

library("foreach")
library("asreml")
```

```
AgeAtMatGenes = c("VGLL3a","AKAP11a","SIX6a") # short version

Cor.All = NULL
Cor.All =
foreach( i = 1:length(AgeAtMatGenes), .combine = "rbind",
.packages=c("asreml","doParallel")) %do% {
        temp2 =
        foreach( j = (1:length(selection))[-which(selection ==
AgeAtMatGenes[i])], .combine = "rbind", .packages=c("asreml","doParallel")) %do%
{

                temp = NULL
                temp = tempDat[ , names(tempDat) == "Stage" |
                                names(tempDat) == "Sample.ID" |
                                names(tempDat) == eval(AgeAtMatGenes[i]) |
                                names(tempDat) == eval(selection[j])]
                names(temp)[3] = "Transcript.1"
                names(temp)[4] = "Transcript.2"
                head(temp)

                model.bivar = NULL
                model.bivar = asreml( cbind(Transcript.1,Transcript.2) ~
at(trait) + at(trait):Stage,
                        rcov = ~ units:us(trait),
                        trace=FALSE,
                        maxiter = 1000,
                        data = temp)

                model.bivar.test = NULL
                model.bivar.test = asreml( cbind(Transcript.1,Transcript.2) ~
at(trait) + at(trait):Stage,
                        rcov = ~ units:diag(trait),
                        trace=FALSE,
                        maxiter = 1000,
                        data = temp)

                cor.test.LRT = if(model.bivar$last.message ==  "LogLikelihood
Converged" & model.bivar$last.message == "LogLikelihood Converged")
LRT.asreml(model.bivar,model.bivar.test) else data.frame(LRT.chisquare = NA, DF
= NA, P.value = NA)
                cor.result =  if(model.bivar$last.message ==  "LogLikelihood
Converged") pin(model.bivar,cor.transcripts ~ V3/sqrt(V2*V4)) else
data.frame(Estimate = NA, SE = NA)

                temp = NULL
                temp = data.frame(Transcript01 = eval(AgeAtMatGenes[i]),
Transcript02 = eval(selection[j]),
                        cor = cor.result$Estimate,
                        cor.se = cor.result$SE,
                        DF = cor.test.LRT$DF,
                        LRT.chisquare = cor.test.LRT$LRT.chisquare,
                        P = cor.test.LRT$P.value)
                rownames(temp) = NULL
```

```
                print(paste(i,"-",j))
                temp
                }
        temp2
        }
head(Cor.All)

# add extra cor of special interest: RD3L - YAP1
temp = NULL
                temp = tempDat[ , names(tempDat) == "Stage" |
                                names(tempDat) == "Sample.ID" |
                                names(tempDat) == "RD3L" |
                                names(tempDat) == "YAP1"]
                names(temp)[3] = "Transcript.1"
                names(temp)[4] = "Transcript.2"
                head(temp)

                model.bivar = NULL
                model.bivar = asreml( cbind(Transcript.1,Transcript.2) ~ 
at(trait) + at(trait):Stage,
                        rcov = ~ units:us(trait),
                        trace=FALSE,
                        maxiter = 1000,
                        data = temp)

                model.bivar.test = NULL
                model.bivar.test = asreml( cbind(Transcript.1,Transcript.2) ~ 
at(trait) + at(trait):Stage,
                        rcov = ~ units:diag(trait),
                        trace=FALSE,
                        maxiter = 1000,
                        data = temp)

                cor.test.LRT = if(model.bivar$last.message ==  "LogLikelihood 
Converged" & model.bivar$last.message == "LogLikelihood Converged") 
LRT.asreml(model.bivar,model.bivar.test) else data.frame(LRT.chisquare = NA, DF 
= NA, P.value = NA)
                cor.result =  if(model.bivar$last.message ==  "LogLikelihood 
Converged") pin(model.bivar,cor.transcripts ~ V3/sqrt(V2*V4)) else 
data.frame(Estimate = NA, SE = NA)

                temp = NULL
                temp = data.frame(Transcript01 = "RD3L", Transcript02 = "YAP1",
                        cor = cor.result$Estimate,
                        cor.se = cor.result$SE,
                        DF = cor.test.LRT$DF,
                        LRT.chisquare = cor.test.LRT$LRT.chisquare,
                        P = cor.test.LRT$P.value)
                rownames(temp) = NULL
Cor.All = rbind(Cor.All,temp)
dim(Cor.All)
# [1] 508   7

any(is.na(Cor.All$cor))
```

```
length(which(is.na(Cor.All$cor)))
# 7 NA

Cor.All[which(is.na(Cor.All$cor)),]
# following genes did not yield model results with any of the three
age-at-maturity genes:
# COL1A1a
# COL1A2
# # TEAD3a - only with SIX6a

Cor.All$P.fine =  1-pchisq(Cor.All$LRT.chisquare, Cor.All$DF)
nrow( Cor.All[Cor.All$P.fine < 0.01,] )
# [1] 26

Cor.All =  Cor.All[which(Cor.All$P.fine < 0.01),]
Cor.All$lower = Cor.All$cor - 2*Cor.All$cor.se
Cor.All$upper = Cor.All$cor + 2*Cor.All$cor.se
range(Cor.All$P.fine)

dim(Cor.All)
# [1] 26 12

write.table(Cor.All,paste0(SubmissionPath,"Significant
correlations.csv"),col.names=TRUE,row.names=FALSE, sep=",")


########################################################
####
#### pairwise comparisons per gene across developmental time points
####
#######################################################

#### subset main data frame for correlating genes
# list with relevant genes
Cor.All$Transcript01 = as.character(Cor.All$Transcript01)
Cor.All$Transcript02 = as.character(Cor.All$Transcript02)
Cor.All.GENES = unique(c(unique(Cor.All$Transcript01),
unique(Cor.All$Transcript02)))
Cor.All.GENES
# [1] "VGLL3a"   "AKAP11a"  "SIX6a"    "RD3L"     "ARHGAP6e" "YAP1"
# [7] "FOXP3a"   "LATS1a"   "NR1I2"    "SOX9d"    "AESb"     "AESc"
#[13] "CYP26B1b" "EGR1d"    "OTX2b"    "PRKAR1Aa" "PRKAR1Bb" "PRKAR2Ad"
#[19] "RTN1"    "SIX3b"    "SLC38A6"  "STK3a"    "TEAD1b"   "VDRAb"

# subset
str(RNADataFrame.detected)

"RNADataFrame.detected.SUB" = NULL
length(unique(RNADataFrame.detected$Transcript.ID))
# [1] 170
dim(RNADataFrame.detected)
# [1] 3060    8
RNADataFrame$Transcript.ID = as.character(RNADataFrame$Transcript.ID)
RNADataFrame.SUB = RNADataFrame[which( RNADataFrame$Transcript.ID %in%
```

```
Cor.All.GENES ),]
length(unique(RNADataFrame.SUB$Transcript.ID))
# [1] 24

RNADataFrame.detected.SUB = RNADataFrame.detected[which(
RNADataFrame.detected$Transcript.ID %in% Cor.All.GENES ),]
dim(RNADataFrame.detected.SUB)
# [1] 432   7
length(unique(RNADataFrame.detected.SUB$Transcript.ID))
# [1] 24

# run model
library("asreml")

RNADataFrame.detected.SUB$Transcript.ID =
as.character(RNADataFrame.detected.SUB$Transcript.ID)
RNADataFrame.detected.SUB =
RNADataFrame.detected.SUB[order(RNADataFrame.detected.SUB$Transcript.ID),]

# simple model with common residual variance
TempGeneDiff = asreml(values ~ 1 + Transcript.ID*Stage,
        random = ~ Sample.ID,
        rcov = ~ units,
        data = RNADataFrame.detected.SUB)
TempGeneDiff$loglik
# [1] -1622.664

# heteroscedastic residual var:
dotplot(residuals(TempGeneDiff) ~ RNADataFrame.detected.SUB$Transcript.ID,
        scales = list(x=list(rot=45)))

RNADataFrame.detected.SUB$hom.residuals = residuals(TempGeneDiff)

round(tapply(RNADataFrame.detected.SUB$hom.residuals,RNADataFrame.detected.SUB$T
ranscript.ID,var))
#    AESb     AESc  AKAP11a ARHGAP6e CYP26B1b    EGR1d   FOXP3a   LATS1a
#    2932      919      240      192      229      382      260      354
#   NR1I2    OTX2b PRKAR1Aa PRKAR1Bb PRKAR2Ad     RD3L     RTN1    SIX3b
#     272       97      482      470      355      224    57803       92
#   SIX6a  SLC38A6    SOX9d    STK3a   TEAD1b    VDRAb   VGLL3a     YAP1
#     125      238      951      653      465      470      242      277
RNADataFrame.detected.SUB$hom.residuals = NULL

# re-fit to account for heteroscedastic residuals var
TempGeneDiff.HET = asreml(values ~ 1 + Transcript.ID*Stage,
        random = ~ Sample.ID,
        rcov = ~ at(Transcript.ID):units,
        data = RNADataFrame.detected.SUB)
TempGeneDiff.HET$loglik
# [1] -1207.33

# formally test whether model is better
LRT = 2*(TempGeneDiff.HET$loglik - TempGeneDiff$loglik)
round(LRT,1)
```

```
# [1] 830.7
1-pchisq(LRT,length(unique(RNADataFrame.detected.SUB$Transcript.ID))-1)
# [1] 0

# check model fit - quite perfect
residual.outlier.plot.asreml( TempGeneDiff.HET )
random.outlier.plot.asreml( TempGeneDiff.HET )

# variances
varcomp.nice(TempGeneDiff.HET)
#                                   component  std.error z.ratio   V
# Sample.ID!Sample.ID.var              6.6989     4.0050    1.67  V1
# Transcript.ID_AESb!variance      5170.4137  1955.4953    2.64  V2
# Transcript.ID_AESc!variance      1739.8864   658.7062    2.64  V3
# Transcript.ID_AKAP11a!variance     49.9040    19.8613    2.51  V4
# Transcript.ID_ARHGAP6e!variance    17.0175     7.5663    2.25  V5
# Transcript.ID_CYP26B1b!variance   350.8055   133.6689    2.62  V6
# Transcript.ID_EGR1d!variance      106.4250    41.2760    2.58  V7
# Transcript.ID_FOXP3a!variance      65.5454    25.7273    2.55  V8
# Transcript.ID_LATS1a!variance     252.3176    96.2892    2.62  V9
# Transcript.ID_NR1I2!variance      293.1825   111.8461    2.62 V10
# Transcript.ID_OTX2b!variance      264.9650   101.2800    2.62 V11
# Transcript.ID_PRKAR1Aa!variance   958.5806   363.5429    2.64 V12
# Transcript.ID_PRKAR1Bb!variance  1018.1086   385.8621    2.64 V13
# Transcript.ID_PRKAR2Ad!variance   682.8654   259.2513    2.63 V14
# Transcript.ID_RD3L!variance        33.6133    13.7879    2.44 V15
# Transcript.ID_RTN1!variance    77506.6386 29296.1835    2.65 V16
# Transcript.ID_SIX3b!variance      145.6651    56.2236    2.59 V17
# Transcript.ID_SIX6a!variance       11.9765     5.7760    2.07 V18
# Transcript.ID_SLC38A6!variance    173.5972    66.6156    2.61 V19
# Transcript.ID_SOX9d!variance     1313.7821   497.4583    2.64 V20
# Transcript.ID_STK3a!variance     1497.1069   567.0328    2.64 V21
# Transcript.ID_TEAD1b!variance     709.8276   269.3990    2.63 V22
# Transcript.ID_VDRAb!variance      220.6548    84.4731    2.61 V23
# Transcript.ID_VGLL3a!variance      44.1570    17.7572    2.49 V24
# Transcript.ID_YAP1!variance       148.0144    56.8997    2.60 V25

# test model terms
wald.nice(TempGeneDiff.HET)
#                      Df denDF   F.con Margin Pr sign
# (Intercept)           1  13.4 4810.00      -  0  ***
# Transcript.ID        23  95.2  877.50      A  0  ***
# Stage                 3  13.4   31.41      A  0  ***
# Transcript.ID:Stage 69 140.9   25.78      B  0  ***

# predict means and se of differences from model
predict.TempGeneDiff.HET = predict(TempGeneDiff.HET,
        classify = "Transcript.ID:Stage",
        sed = TRUE)$prediction
predict.TempGeneDiff.HET.pvals = predict.TempGeneDiff.HET$pvals
predict.TempGeneDiff.HET.pvals$rowname = 1:nrow(predict.TempGeneDiff.HET.pvals)
head(predict.TempGeneDiff.HET.pvals)

predict.TempGeneDiff.HET.sed = predict.TempGeneDiff.HET$sed
```

```
predict.TempGeneDiff.HET.sed

# pairwise comparisons
Comp.All = NULL
for(Transcript in
1:length(unique(predict.TempGeneDiff.HET.pvals$Transcript.ID)))
{
        SELECT = which( predict.TempGeneDiff.HET.pvals$Transcript.ID ==
unique(predict.TempGeneDiff.HET.pvals$Transcript.ID)[Transcript])
        temp = NULL
        temp = predict.TempGeneDiff.HET.pvals[SELECT,]
        sed = predict.TempGeneDiff.HET.sed

        Comp.temp =
        data.frame(
                Transcript.ID =
rep(unique(predict.TempGeneDiff.HET.pvals$Transcript.ID)[Transcript],6),
                Contrast = c(   "1-2","1-3","1-4","2-3","2-4","3-4" ),
                Diff = c(       temp[1,]$predicted.value -
temp[2,]$predicted.value,
                                        temp[1,]$predicted.value -
temp[3,]$predicted.value,
                                        temp[1,]$predicted.value -
temp[4,]$predicted.value,
                                        temp[2,]$predicted.value -
temp[3,]$predicted.value,
                                        temp[2,]$predicted.value -
temp[4,]$predicted.value,
                                        temp[3,]$predicted.value -
temp[4,]$predicted.value ),
                SED = c(        sed[temp[1,]$rowname,temp[2,]$rowname],
                                        sed[temp[1,]$rowname,temp[3,]$rowname],
                                        sed[temp[1,]$rowname,temp[4,]$rowname],
                                        sed[temp[2,]$rowname,temp[3,]$rowname],
                                        sed[temp[2,]$rowname,temp[4,]$rowname],
                                        sed[temp[3,]$rowname,temp[4,]$rowname] )
)
                Comp.All = rbind(Comp.All,Comp.temp)
        }
Comp.All

Comp.All$t = Comp.All$Diff / Comp.All$SED
Comp.All$P = 2*(1-pt(abs(Comp.All$t), df = 13.4))
Comp.All$FDR = p.adjust(Comp.All$P,"BH")
Comp.All$Flag = ifelse(Comp.All$FDR < 0.05, "*",
                                ifelse(Comp.All$FDR < 0.01, "**",
                                ifelse(Comp.All$FDR < 0.001, "***", "")))

# list with significant genes
temp = NULL
temp = as.character( unique(Comp.All[Comp.All$Flag !="",]$Transcript.ID) )
temp
#  [1] "AESb"     "AESc"     "AKAP11a"  "ARHGAP6e" "CYP26B1b" "EGR1d"
#  [7] "FOXP3a"   "LATS1a"   "OTX2b"    "PRKAR1Aa" "PRKAR1Bb" "PRKAR2Ad"
```

```
# [13] "RD3L"     "SIX3b"    "SIX6a"    "SLC38A6"  "SOX9d"    "STK3a"
# [19] "TEAD1b"   "VDRAb"    "VGLL3a"   "YAP1"

# list with non-significant genes
temp2 = as.character(unique(Comp.All[Comp.All$Flag == "",]$Transcript.ID))
temp2
#  [1] "AESb"     "AESc"     "AKAP11a"  "ARHGAP6e" "CYP26B1b" "EGR1d"
#  [7] "FOXP3a"   "LATS1a"   "NR1I2"    "PRKAR1Aa" "PRKAR1Bb" "PRKAR2Ad"
# [13] "RD3L"     "RTN1"    "SIX3b"    "SIX6a"    "SLC38A6"  "TEAD1b"
# [19] "VDRAb"    "VGLL3a"   "YAP1"

# correlated genes that are not different across developmental
temp2[which(!temp2 %in% temp)]
# [1] "NR1I2" "RTN1"

# make correlation groups -
VGLL3a.Genes = Cor.All[which(Cor.All$Transcript01 == "VGLL3a"),]$Transcript02
AKAP11a.Genes = Cor.All[which(Cor.All$Transcript01 == "AKAP11a"),]$Transcript02
SIX6a.Genes = Cor.All[which(Cor.All$Transcript01 == "SIX6a"),]$Transcript02
VGLL3a.Genes
AKAP11a.Genes
SIX6a.Genes

predict.TempGeneDiff.HET.pvals$Transcript.ID =
as.character(predict.TempGeneDiff.HET.pvals$Transcript.ID)

predict.TempGeneDiff.HET.pvals$VGLL3a.Group = NA
predict.TempGeneDiff.HET.pvals$VGLL3a.Group[which(predict.TempGeneDiff.HET.pvals
$Transcript.ID %in% c(VGLL3a.Genes,"VGLL3a"))] = "yes"
predict.TempGeneDiff.HET.pvals$AKAP11a.Group = NA
predict.TempGeneDiff.HET.pvals$AKAP11a.Group[which(predict.TempGeneDiff.HET.pval
s$Transcript.ID %in% c(AKAP11a.Genes,"AKAP11a"))] = "yes"
predict.TempGeneDiff.HET.pvals$SIX6a.Group = NA
predict.TempGeneDiff.HET.pvals$SIX6a.Group[which(predict.TempGeneDiff.HET.pvals$
Transcript.ID %in% c(SIX6a.Genes,"SIX6a"))] = "yes"

xyplot( (predicted.value) ~ Stage  ,
        groups = Transcript.ID, auto.key=list(columns=6),
        main = "VGLL3a.Group",
        type=c("p","l"),
        scales = list(relation="free"),
        data = droplevels(predict.TempGeneDiff.HET.pvals[which(
                predict.TempGeneDiff.HET.pvals$Transcript.ID %in% temp &
predict.TempGeneDiff.HET.pvals$VGLL3a.Group == "yes"),]))

xyplot( (predicted.value) ~ Stage  ,
        groups = Transcript.ID, auto.key=list(columns=6),
        main = "SIX6a.Group",
        type=c("p","l"),
        scales = list(relation="free"),
        data = droplevels(predict.TempGeneDiff.HET.pvals[which(
                predict.TempGeneDiff.HET.pvals$Transcript.ID %in% temp &
predict.TempGeneDiff.HET.pvals$SIX6a.Group == "yes"),]))
```

```
xyplot( (predicted.value) ~ Stage  ,
        groups = Transcript.ID, auto.key=list(columns=6),
        main = "AKAP11a.Group",
        type=c("p","l"),
        scales = list(relation="free"),
        data = droplevels(predict.TempGeneDiff.HET.pvals[which(
                predict.TempGeneDiff.HET.pvals$Transcript.ID %in% temp &
predict.TempGeneDiff.HET.pvals$AKAP11a.Group == "yes"),]))

Comp.All$Contrast = paste0("Contrast ",Comp.All$Contrast)

path = "D:/Documents/Research Projects/2018 Johanna Correlation/Data"

write.table(Comp.All,paste0(SubmissionPath,"Developmental Stages -
Comparisons.csv"),col.names=TRUE,row.names=FALSE, sep=",")
write.table(predict.TempGeneDiff.HET.pvals,paste0(SubmissionPath,"Developmental
Stages - Means.csv"),col.names=TRUE,row.names=FALSE, sep=",")
```
